# Causal effects of lifetime smoking on risk for depression and schizophrenia: Evidence from a Mendelian randomisation study

**DOI:** 10.1101/381301

**Authors:** Robyn E Wootton, Rebecca C Richmond, Bobby G Stuijfzand, Rebecca B Lawn, Hannah M Sallis, Gemma M. J. Taylor, Gibran Hemani, Hannah J. Jones, Stanley Zammit, George Davey Smith, Marcus R Munafò

**Affiliations:** School of Experimental Psychology, University of Bristol, Bristol, BS8 1TU; Department of Population Health Sciences, Bristol Medical School, University of Bristol, Bristol, BS8 2PR; MRC Integrative Epidemiology Unit, University of Bristol, Bristol, BS8 2PR; NIHR Biomedical Research Centre at the University Hospitals Bristol NHS Foundation Trust and the University of Bristol, (BS8 2BN).; UK Centre for Tobacco and Alcohol Studies, University of Bristol, Bristol, BS8 1TU; Jean Golding Institute, Royal Fort House, University of Bristol, Bristol, BS8 1UH; MRC Centre for Neuropsychiatric Genetics and Genomics, Division of Psychological Medicine and Clinical Neurosciences, Cardiff University, Cardiff CF24 4HQ

## Abstract

**Background:** Smoking prevalence is higher amongst individuals with schizophrenia and depression compared to the general population. Mendelian randomisation (MR) can examine whether this association is causal using genetic variants identified in genome-wide association studies (GWAS).

**Methods:** We conducted a GWAS of lifetime smoking behaviour (capturing smoking duration, heaviness and cessation) in a sample of 462,690 individuals from the UK Biobank, and validated the findings via two-sample MR analyses of positive control outcomes (e.g., lung cancer). Having established the validity of our instrument, we used bi-directional two-sample Mendelian randomisation to explore its effects on schizophrenia and depression.

**Outcomes:** There was strong evidence to suggest smoking is a causal risk factor for both schizophrenia (OR = 2.27, 95% CI = 1.67 - 3.08, P < 0.001) and depression (OR = 1.99, 95% CI = 1.71 - 2.32, P < 0.001). We also found some evidence that genetic risk for both schizophrenia and depression cause increased lifetime smoking (β = 0.022, 95% CI = 0.005 - 0.038, P = 0.009; β= 0.091, 95% CI = 0.027 - 0.155, P = 0.005).

**Interpretation:** These findings suggest that the association between smoking, schizophrenia and depression is due, at least in part, to a causal effect of smoking, providing further evidence for the detrimental consequences of smoking for mental health.

**Funding:** This work was supported by the Medical Research Council Integrative Epidemiology Unit, the NIHR Biomedical Research Centre, University Hospitals Bristol NHS Foundation Trust and the University of Bristol.

**Research in context:** *Evidence before this study:* The association between smoking and mental health (especially schizophrenia and depression) is often assumed to be the result of self-medication (for example, to alleviate symptoms). However, more recent evidence has suggested that smoking might also be a risk factor for schizophrenia and depression. This alternative direction of effect is supported by meta-analyses and previous prospective observational evidence using related individuals to control for genetic and environmental confounding. However, observational evidence cannot completely account for confounding or the possibility of reverse causation. One way to get around these problems is Mendelian randomisation (MR). Previous MR studies of smoking and mental health have not shown an effect of smoking on depression and are inconclusive for the effects of smoking on schizophrenia. However, these studies have only looked at individual aspects of smoking behaviour and some studies required stratifying participants into smokers and non-smokers, reducing power.

*Added value of this study:* We have developed a novel genetic instrument for lifetime smoking exposure which can be used within a two-sample MR framework, using publicly-available GWAS summary statistics. We were therefore able to test the bi-directional association between smoking with schizophrenia and depression to see if the effects are causal. We found strong evidence to suggest that smoking is a causal risk factor for both schizophrenia and depression. There was some evidence to suggest that risk of schizophrenia and depression increases lifetime smoking (consistent with the self-medication hypothesis) but the effects were stronger for depression than schizophrenia.

*Implications of all the available evidence:* This study was the first to demonstrate evidence for an effect of lifetime smoking exposure on risk of schizophrenia and depression within a causal inference framework. This emphasises the detrimental public health consequences of smoking, not just for physical health, but also to mental illness.

## Introduction

Smoking is a major risk factor for lung cancer, cardiovascular and respiratory diseases making it the leading cause of preventable death worldwide^1^. In developed nations, smoking is more common amongst individuals with mental health conditions^2^–^5^, in particular schizophrenia^6^ and depression^7–9^. In the UK, estimates suggest that up to 45% of individuals with schizophrenia, and 31% of individuals with depression smoke^6^, compared to around 15% of the general population^10^. Individuals with mental health conditions smoke more heavily^2^ and experience up to 18 years reduced life expectancy compared with the general population^6,11^. Much of this reduction can be explained by smoking related diseases^6^, making it important to understand the relationship between smoking and mental health.

It is often assumed that the association between mental health and smoking can be explained by a self-medication model – that is, symptoms of mental illness, or side effects of psychiatric medications, are alleviated by the chemical properties of tobacco^12–15^. However, observational evidence cannot determine whether the association between smoking and mental health is causal or the result of confounding^16^. Furthermore, traditional observational evidence cannot robustly identify the direction of causation^16^, and there is growing evidence to suggest that smoking may be a causal risk factor for poor mental health. The genome-wide association study (GWAS) of schizophrenia conducted by the Psychiatric Genomics Consortium (PGC) found that variants in the gene cluster *CHRNA5-A3-B4* were associated with increased schizophrenia risk^17^. These variants are known to be strongly associated with heaviness of smoking^18–21^. Therefore, one interpretation is a causal effect of smoking on schizophrenia^22^. Furthermore, there is evidence of a genetic correlation between smoking and schizophrenia^23^ and polygenic risk scores for schizophrenia associate with smoking status warranting further investigation of possible causal effects^24^. Prospective observational studies using related individuals to control for genetic and environmental confounding have also suggested a dose-response effect of smoking on schizophrenia^25^ and depression^26^. Meta-analyses show further evidence for an increased relative risk of schizophrenia in smokers over non-smokers^27,28^ and a reduction in depressive symptoms following smoking cessation^29^. Although these studies suggest a potential causal effect, more robust methods are required to triangulate evidence and allow for stronger causal inference^30^.

One way to overcome the limitations of residual confounding and reverse causation is Mendelian randomisation (MR)^31,32^. This method uses genetic variants to proxy for an exposure in an instrumental variable analysis to estimate the causal effect on an outcome^16^. Previous MR studies have failed to show any clear evidence for a causal effect of smoking on depression^33–35^ and show suggestive but inconclusive evidence for an effect of smoking on schizophrenia^35,36^. However, the genetic instruments for smoking used in these MR studies are limited, only capturing individual aspects of smoking behaviour such as smoking initiation or smoking heaviness, rather than total lifetime smoking exposure^33–36^. Furthermore, any instrument for smoking heaviness requires stratifying samples into smokers and non-smokers. Stratification is not possible using the most common MR method; two-sample MR. In two-sample MR, SNP-exposure and SNP-outcome effects are estimated in two independent samples and the effect sizes are obtained from GWAS summary statistics. Therefore, in this context stratification is not possible because GWAS summary data do not provide individual level data regarding smoking status.

Here we report the development of a novel genetic instrument for lifetime smoking exposure that takes into account smoking status (i.e., ever and never smokers), and among ever smokers takes into account smoking duration, heaviness and cessation. This instrument can therefore be used in unstratified samples of smokers and non-smokers. We validate this instrument in an independent sample and against positive control disease outcomes (lung cancer, coronary artery disease (CAD) and DNA methylation). We next apply the instrument in MR analyses to determine whether the observational associations between smoking, schizophrenia and depression are causal, and the directionality of these relationships.

## Methods

### Instrument discovery for lifetime smoking index

#### Sample

We generated the lifetime smoking measure using data from the UK Biobank, a national health research resource of 502,647 participants aged 40-69 years, recruited from across the United Kingdom between 2006 and 2010 (http://www.ukbiobank.ac.uk). Our sample consisted of 462,690 individuals of European ancestry who had phenotype data and passed genotype inclusion criteria (54% female; mean age = 56.7 years; SD = 8.0 years). Overall, 30% of the sample had ever smoked (8% current smokers and 22% former smokers).

#### Measures of smoking

Smoking measures available in the UK Biobank were self-reported and collected at initial assessment. They included: smoking status (current, former, never – field 20116), age at initiation in years (fields 3436 and 2867), age at cessation in years (field 2897) and number of cigarettes smoked per day (fields 3456 and 2887). Anyone self-reporting to smoke more than 100 cigarettes per day was contacted to confirm their response. Hand-rolled cigarette smokers were told one gram of tobacco equates to one cigarette. We calculated duration of smoking and time since cessation. Individuals reporting smoking fewer than 1 cigarette a day or more than 150 cigarettes a day were excluded.

#### Construction of the lifetime smoking index

Following the method outlined by Leffondré and colleagues^37^, we combined the smoking measures into a lifetime smoking index along with a simulated half-life (τ) constant. Half-life captures the exponentially decreasing effect of smoking at a given time on health outcomes. The value of half-life was determined by simulating the effects of lifetime smoking on lung cancer and overall mortality in the UK Biobank. Both suggested the best fitting value was 18. For full details on construction of the lifetime smoking instrument see Supplementary Note.

#### Genome-wide association study of lifetime smoking index

For full details of genotyping and exclusion procedures see Supplementary note. GWAS was conducted using the UK Biobank GWAS pipeline set up for the MRC Integrative Epidemiology Unit^38^. BOLT-LMM was used to conduct the analysis^39^, which accounts for population stratification and relatedness using linear mixed modelling. Genotyping chip and sex were included as covariates. As a sensitivity analysis, we reran the GWAS without controlling for genotype chip because the UK BiLEVE sub-sample (for which a different genotyping chip was used) were selected on the basis of smoking status. Genome-wide significant SNPs were selected at P < 5 × 10^-8^ and were clumped to ensure independence at linkage disequilibrium (LD) r^2^ = 0.001 and a distance of 10,000 kb using the clump_data command in the TwoSampleMR package^40^ which uses PLINK software^41^ and only clumps SNPs within the specified kb window.

#### Prediction in an independent sample

To validate our instrument, we used the Avon Longitudinal Study of Parents and Children (ALSPAC). For details of this cohort and measures, see Supplementary Note. PLINK was used to generate a polygenic risk score in ALSPAC^41^. Linear regression was used to estimate the percentage variance of lifetime smoking explained by the polygenic score.

### Instrument validation

#### Positive control outcomes

We tested our genetic instrument using the positive controls of lung cancer, CAD and hypomethylation at the aryl-hydrocarbon receptor repressor (*AHRR*) site cg05575921 because smoking has been robustly shown to predict these outcomes^1,42^. We conducted these analyses using GWAS summary data in a two-sample MR framework using our exposure instrument for lifetime smoking from our GWAS in the UK Biobank. For details of the outcome GWAS see Supplementary Note.

#### Two-sample Mendelian randomisation of positive controls

Analyses were conducted using MR Base, an R^43^ package for two-sample MR^40^. We compared results across five different MR methods: inverse-variance weighted, MR Egger^44^, weighted median^45^, weighted mode^46^ and MR RAPS^47^. Each method makes different assumptions and therefore a consistent effect across multiple methods strengthens causal evidence^48^. If a SNP was unavailable in the outcome GWAS summary statistics, then proxy SNPs were searched for with a minimum LD r^2^ = 0.8. We aligned palindromic SNPs with MAF<0.3.

### Bidirectional two-sample Mendelian randomisation of mental health

#### Data sources

For lifetime smoking, we used the summary data from our GWAS of lifetime smoking in the UK Biobank. For schizophrenia, we used summary data from the PGC consortium GWAS, which comprises 36,989 cases and 113,075 controls of mixed ancestry^17^. Cases were a combination of individuals with schizophrenia and schizoaffective disorder mostly diagnosed by clinicians but some samples used research-based assessment. Post hoc analyses showed that ascertainment method did not affect results^17^. A sensitivity analysis was performed using the GWAS summary data meta-analysed with a further 11,260 cases and 24,542 controls^49^. For depression, we used the most recent GWAS summary data from the PGC for major depression, which comprises 130,664 major depression cases and 330,470 controls of European ancestry^50^. Cases were either diagnosed with major depressive disorder (MDD) on inpatient or medical health records or self-reported having a diagnosis or treatment for depression. Therefore, the authors use the term major depression over diagnosed MDD^50^. 23andMe data (75,607 cases and 231,747 controls) were excluded when major depression was the outcome because genome-wide summary statistics are not available with 23andMe data included.

#### Genetic instruments

For lifetime smoking, we identified 126 independent loci at genome-wide significance, which explained 0.36% of the variance. The genetic instrument for schizophrenia came from the PGC GWAS, which identified 114 independent SNPs at 108 loci explaining around 3.4% of the variance in schizophrenia liability^17^. Finally, the genetic instrument for major depression from the PGC GWAS was 40 genome-wide significant SNPs which explain 1.9% of the variance in liability^50^. As a further sensitivity analysis, we compared our lifetime smoking instrument with the most recent GWAS of smoking initiation from GSCAN^51^. They identified 378 conditionally independent genome-wide significant SNPs, which explain 2% of the variance in smoking initiation^51^.

#### Two-sample Mendelian randomisation of mental health outcomes

Analyses were run using the MR Base R Package^40,43^ and compared across the five different methods. This time, analyses of GWAS summary data were run bi-directionally, with lifetime smoking first as the exposure and then as the outcome. Steiger filtering was conducted to confirm the direction of effect^52^. If a SNP from the instrument was unavailable in the outcome, an attempt to find proxies was made with a minimum LD r^2^ = 0.8 and palindromic SNPs were aligned with MAF<0.3.

## Results

### Lifetime smoking score construction

After excluding individuals who did not pass genotype exclusions and who had missing smoking phenotype data, 462,690 individuals remained for the GWAS. Of these individuals, 249,318 were never smokers (54%), 164,649 were previous smokers (36%) and 48,723 (11%) were current smokers. The mean value of lifetime smoking score was 0.359 (SD=0.694). A standard deviation increase in lifetime smoking score is equivalent to an individual smoking 20 cigarettes a day for 15 years and stopping 17 years ago or an individual smoking 60 cigarettes a day for 13 years and stopping 22 years ago.

### Instrument discovery for lifetime smoking index

The results of our GWAS of lifetime smoking (N = 462,690) are presented in Figure 1 and Supplementary Figure S1. The most strongly associated regions on chromosome 15 and chromosome 9 have been previously shown to be associated with smoking heaviness and cessation respectively^53^. We identified 10,415 SNPs at the genome-wide level of significance (P < 5 × 10^-8^). After clumping and filtering, 126 independent SNPs remained. A full list of these SNPs and their z-scored effect sizes can be found in Supplementary Table S1. In the ALSPAC independent sample, the 126 SNPs explained 0.36% of the variance in lifetime smoking (P = 0.002).

**Figure 1.**
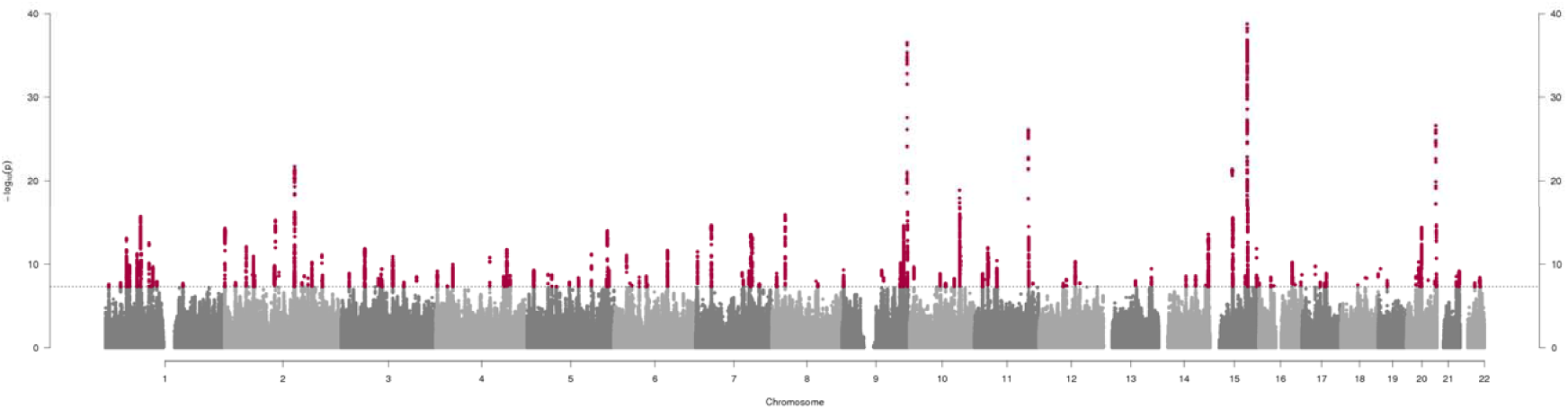
Manhattan plot of genome-wide association study lifetime smoking index (N = 462,690). The x-axis represents chromosomal position and the y-axis represents −log_10_ P-value for the association of each single nucleotide polymorphism (SNP) with the lifetime smoking index using an additive model and linear regression. The dashed line indicates the genome-wide level of significance (P < 5 × 10^-8^) and genome-wide significant SNPs are indicated in red.

### Instrument validation

#### Two-sample Mendelian randomisation of positive controls

We validated the genetic instrument for lifetime smoking exposure using two-sample MR of smoking on positive control outcomes: lung cancer, CAD and hypomethylation at the *AHRR* locus. All five MR methods indicated the expected direction of effect (see Table 1) increasing risk of disease outcomes and decreasing *AHRR* methylation, with the exception of the MR Egger SIMEX adjusted estimates for CAD. However, these should be interpreted with caution given the low I^2^ _GX_ (see Supplementary Table S2)^54^. There was strong evidence of an effect of lifetime smoking on increased odds of lung cancer and CAD. There was weaker evidence of an effect on AHRR methylation, possibly due to lower power. Sensitivity analyses are presented in Supplementary Figures S2-S7. There was evidence of significant heterogeneity in the SNP-exposure effects (see Supplementary Table S3); however, tests of MR Egger intercepts generally indicated weak evidence of directional pleiotropy (see Supplementary Table S4), with the exception of CAD.

**Table 1.** **Two-sample Mendelian randomisation analyses of the effect of lifetime smoking on coronary artery disease, lung cancer and *AHRR* Methylation.**

| **Outcome** | **Method** | **OR (95% CI)** | **P-value** |
| --- | --- | --- | --- |
| **Coronary Artery Disease** | Inverse-Variance Weighted | 1.56 (1.34, 1.82) | $1.19 \times 10^{-08}$ |
|  | MR Egger (SIMEX) | 0.76 (0.44, 1.34) | 0.35 |
| | Weighted Median | 1.65 (1.36, 2.00) | $5.40 \times 10^{-07}$ |
|  | Weighted Mode | 1.79 (1.06, 3.03) | 0.03 |
| | MR RAPS | 1.63 (1.40, 1.90) | $5.65 \times 10^{-10}$ |
| **Lung Cancer** | Inverse-Variance Weighted | 4.21 (2.98, 5.96) | $3.49 \times 10^{-16}$ |
| | MR Egger (SIMEX) | 16.64 (3.88, 71.42) | $9.61 \times 10^{-05}$ |
| | Weighted Median | 2.77 (1.91, 4.03) | $8.88 \times 10^{-08}$ |
|  | Weighted Mode | 6.19 (2.07, 18.54) | 0.001 |
| | MR RAPS | 3.71 (2.75, 5.00) | $8.65 \times 10^{-18}$ |
|  |  | **Beta (95% CI)** | **P-value** |
| ***AHRR* Methylation** | Inverse-Variance Weighted | -0.098 (-0.168, -0.028) | 0.006 |
|  | MR Egger (SIMEX) | -0.176 (-0.443, 0.102) | 0.217 |
|  | Weighted Median | -0.125 (-0.228, -0.021) | 0.02 |
|  | Weighted Mode | -0.207 (-0.511, 0.097) | 0.18 |
|  | MR RAPS | -0.095 (-0.171, -0.013) | 0.01 |

### Bidirectional two-sample Mendelian randomisation of mental health

Bi-directional MR analyses provided strong evidence that increased lifetime smoking increases risk of both schizophrenia (OR = 2.27, 95% CI = 1.67 - 3.08, P < 0.001) and depression (OR = 1.99, 95% CI = 1.71 - 2.32, P < 0.001) (see Table 2), with consistent direction of effect across all five MR methods. Again, MR Egger results are the least reliable due to low I^2^_GX_ (see Supplementary Table S2). There was also evidence of a consistent but smaller causal effect of higher genetic risk for schizophrenia on lifetime smoking (β = 0.022, 95% CI = 0.005 - 0.038, P = 0.009) and of genetic risk for depression on lifetime smoking (β= 0.091, 95% CI = 0.027 - 0.155, P = 0.005) (see Table 2). However, the effect was larger and more consistent for depression than for schizophrenia. We saw similar effects using the more recent meta-analysed GWAS for schizophrenia with an additional 11,260 cases (see Supplementary Table S10). The I^2^_GX_ value for schizophrenia and depression was less than 0.6, therefore MR Egger was not conducted (see Supplementary Table S2). There was evidence of significant heterogeneity (see Supplementary Table S3) but MR Egger intercepts suggest directional pleiotropy is not biasing the estimates (see Supplementary Table S4) and Steiger filtering supported the conclusion that the effects operate in both directions (see Supplementary Table S5). A total of 14,260 cases and 15,480 controls from the UK Biobank were included in the latest GWAS for major depression^50^ which could lead to bias from sample overlap^55^. Therefore, this analysis was repeated using summary data from an earlier GWAS of major depression^56^, which showed a consistent direction of effect with weaker statistical evidence, possibly due to reduced sample size (N = 18,759) (see Supplementary Table S6). Bi-directional MR analyses were repeated using the GWAS of lifetime smoking without controlling for genotyping chip and the same pattern of results was observed (see Supplementary Table S7). Finally, analysis of our lifetime smoking instrument was compared with the smoking initiation instrument identified by GSCAN^51^ (see Supplementary Table S8). Results were consistent for both lifetime smoking and smoking initiation as the exposure, with smaller effect sizes for smoking initiation than lifetime smoking. There remained evidence of an effect of depression risk on initiating smoking but even less support of an effect of schizophrenia risk on smoking initiation with inconsistent directions of effect across the various sensitivity measures (see Supplementary Table S8). Further sensitivity tests were conducted and are presented in Supplementary Figures S9-S20.

**Table 2.** **Bidirectional two-sample Mendelian randomisation analyses of the effect of lifetime smoking on schizophrenia and major depression.**

| Exposure | Outcome | Method | OR (95% CI) | P-value |
| --- | --- | --- | --- | --- |
| Smoking | Schizophrenia | Inverse-Variance Weighted | 2.27 (1.67, 3.08) | 1.36 × 10<sup>-07</sup> |
|  |  | MR Egger (SIMEX) | 4.59 (1.49, 14.11) | 0.009 |
|  |  | Weighted median | 2.04 (1.57, 2.64) | 8.21 × 10<sup>-08</sup> |
|  |  | Weighted mode | 1.71 (0.69, 4.23) | 0.25 |
|  |  | MR RAPS | 2.44 (1.84, 3.25) | 9.30 × 10<sup>-10</sup> |
| Smoking | Depression | Inverse-Variance Weighted | 1.99 (1.71, 2.32) | 9.69 × 10<sup>-19</sup> |
|  |  | MR Egger (SIMEX) | 1.09 (0.62, 1.92) | 0.77 |
|  |  | Weighted median | 1.97 (1.65, 2.35) | 3.00 × 10<sup>-14</sup> |
|  |  | Weighted mode | 1.83 (1.19, 2.81) | 0.007 |
|  |  | MR RAPS | 1.99 (1.70, 2.32) | 2.76 × 10<sup>-18</sup> |
|  |  |  | Beta (95% CI) | P-value |
| Schizophrenia | Smoking | Inverse-Variance Weighted | 0.022 (0.005, 0.038) | 0.009 |
|  |  | MR Egger (SIMEX) | - | - |
|  |  | Weighted median | 0.015 (0.004, 0.026) | 0.009 |
|  |  | Weighted Mode | 0.016 (-0.014, 0.045) | 0.31 |
|  |  | MR RAPS | 0.018 (0.003, 0.032) | 0.015 |
| Depression | Smoking | Inverse-Variance Weighted | 0.091 (0.027, 0.155) | 0.005 |
|  |  | MR Egger (SIMEX) | - | - |
|  |  | Weighted median | 0.100 (0.058, 0.141) | 2.77 × 10<sup>-06</sup> |
|  |  | Weighted mode | 0.109 (0.037, 0.182) | 0.005 |
|  |  | MR RAPS | 0.078 (0.014, 0.141) | 0.016 |

## Discussion

We conducted a GWAS of lifetime smoking exposure which provides a novel genetic instrument that can be used in two-sample MR of summary data without the need to stratify on smoking status. Our validation analyses confirm that the lifetime smoking instrument predicts increased risk of lung cancer, CAD and hypomethylation at the *AHRR* locus. Of the 126 independent loci identified, many associations are in novel regions, but we also replicated associations in the *CHRNA5-A3-B4* gene cluster on chromosome 15 previously associated with smoking heaviness and a region near the *DBH* gene on chromosome 9 previously associated with smoking cessation^53^.

We used this novel genetic instrument to explore causal pathways between smoking, schizophrenia and depression. The two-sample MR results provide evidence that smoking is a causal risk factor for both schizophrenia and depression. This supports prospective observational evidence controlling for genetic confounding^25,26^, as well as meta-analyses of observational studies^27,29^ (although it should be noted that these meta-analyses include estimates not adjusted for known confounders, for example, cannabis use). Effect sizes were similar to a more recent meta-analysis that did adjust for multiple confounders and found a two-fold increased risk of schizophrenia in smokers compared with non-smokers^28^. Some studies which adjust for potential confounders find the effect of smoking attenuates to the null^57^ or even becomes protective^58^, demonstrating that there are likely to be substantial confounding effects in observational studies. Previous MR studies have not found clear evidence to support smoking as a causal risk factor for either schizophrenia or depression^33–36^, but our approach offers greater power and better captures lifetime smoking rather than individual components of smoking behaviour and enables two-sample MR analysis using summary data in unstratified samples. However, it is not possible to precisely estimate from our results what proportion of the observational association between smoking, schizophrenia and depression is causal, or the population attributional fraction of these disorders due to smoking.

In support of the self-medication hypothesis^12^, we found evidence to suggest that genetic risk for schizophrenia and depression increases lifetime smoking. This supports previous observational evidence^13–15^ and might explain why smoking rates remain so high amongst individuals with schizophrenia and depression compared with the general population^59^. However, the evidence was stronger for self-medication effects in depression than schizophrenia and when using smoking initiation as the outcome rather than lifetime smoking, effects attenuated to the null. Therefore, maybe any self-medication effects of schizophrenia are only on heaviness and duration of smoking (captured by the lifetime smoking index) rather than initiation. However, it is important to note that the effects might be weaker because MR methods typically capture the long-term effects of exposures^60^. Clinical observations suggest that smoking increases when symptoms are exacerbated among patients with schizophrenia, which is typically interpreted as evidence of self-medication^61^. However, the mechanisms underlying acute effects of psychiatric symptoms on short-term smoking behaviour may differ from the chronic effects of smoking on mental health, with cigarettes having positive effects on symptoms only in the short-term. This highlights the need to differentiate between acute and chronic exposures, and short- and long-term effects, with MR generally being better suited to interrogating chronic exposures and long-term effects.

The effects of schizophrenia and depression on lifetime smoking might be explained by the misattribution hypothesis. This proposes that smokers misattribute the ability of cigarettes to relieve withdrawal, to their ability to relieve symptoms of psychological distress^62,63^. For example, withdrawal symptoms include depressed mood, anxiety and irritability, and smoking a cigarette alleviates those symptoms^64^. Since many withdrawal symptoms are similar to the negative symptoms of schizophrenia and mood symptoms in depression, their alleviation by smoking could give rise to the strong belief that smoking helps to alleviate mental health symptoms. This, in turn, might serve to maintain smoking among individuals with schizophrenia and depression^65^. Given that the symptoms of withdrawal are most similar to the symptoms of depression, this might further explain why we see stronger effects of depression risk on smoking than schizophrenia risk on smoking.

A potential biological mechanism for the bi-directional causal effects of smoking, schizophrenia and depression could be neuroadaptations in the dopaminergic and serotonin systems. Nicotine acts on nicotinic cholinergic receptors in the brain stimulating the release of neurotransmitters including dopamine and serotonin^65^. Dopamine and serotonin dysfunction have been implicated in the aetiology of schizophrenia and depression respectively^66,67^. It is plausible, therefore, that disruption of these pathways has a causal effect on these disorders. Alternatively, cannabis use could be a mediating mechanism for the effects of smoking on schizophrenia and depression. In prospective studies, cigarette smoking has been shown to increase risk of cannabis dependence even after adjusting for cannabis use at baseline^68^. There is strong evidence suggesting that cannabis use increases the risk of psychosis and affective disorders^69^. This vertical pleiotropy does not violate the assumptions of MR, but simply means there are intermediate mechanisms operating between the exposure and the outcome. However, the strong effects we observe for lifetime smoking suggest that any mediating influence of cannabis use is likely to only partially account for these effects, given the relatively low prevalence of cannabis use (e.g., annual prevalence in the UK of ~7% in 2010^70^). Multivariable MR analysis of tobacco and cannabis use would help resolve this question.

Future work should attempt to elucidate the underlying mechanisms with a hope to intervene, inform public health messages or further advance our knowledge on the aetiology of mental illness. In particular, it will be important to consider other constituents of tobacco smoke to determine whether it is exposure to nicotine or some other constituent that increases risk of schizophrenia and depression. This is particularly important in the context of the recent growth in electronic cigarette use. Furthermore, future work should aim to disentangle the effects of maternal smoking during pregnancy from the effects of the offspring’s own smoking behaviour using samples with genetic data on multiple generations.

### Strengths and Limitations

Our study is the first to generate a genetic instrument for lifetime smoking behaviour in a large sample, which allows the use of two-sample MR with summary data from unstratified samples to answer questions about the association between smoking and other health outcomes. However, there are some limitations which should be noted. First, there is evidence to suggest that even after seemingly controlling for population structure in GWAS of samples as large as the UK Biobank, coincident apparent structure remains^71^. This might confound the association between smoking and mental health, increasing the risk of false positives. As independent samples with adequate sample size become available, the influence of structure should be further explored. However, it is reassuring that our instrument predicted lifetime smoking in the ALSPAC replication sample, where such issues would not arise in the same manner.

Second, sample overlap in two-sample MR can bias results towards the observational estimate^55^. There was some sample overlap between the major depression GWAS^50^ and the UK Biobank (used to derive the lifetime smoking instrument) meaning that the effects could be inflated. Therefore, a sensitivity analysis was conducted using a previous GWAS of major depression^56^ which showed the same direction of effect despite lower power. This gives us confidence in the bi-directional effects of smoking and depression, despite sample overlap. This sensitivity analysis also addresses recent concerns over the specificity of the most recent GWAS for major depression^72^. Comparing self-reported “seeking help for depression” with DSM diagnosed MDD yielded different results^72^. However, the earlier GWAS of depression did use DSM diagnosed cases only^56^ and showed the same direction of effect despite lower power.

Third, including multiple aspects of smoking behaviour could have introduced more potential for horizontal pleiotropy. The more diffuse the definition of smoking, the more lifestyle factors might be correlated, making it especially important to test for horizontal pleiotropy. We conducted multiple sensitivity analyses (which all make different and largely uncorrelated assumptions), and all demonstrated the same direction of effect. This increases our confidence that the results are not biased by pleiotropy. Furthermore, MR Egger intercepts did not show evidence of directional pleiotropy for schizophrenia or depression. However, further work should still attempt to understand the biological mechanisms underpinning the association in order to reduce the likelihood of pleiotropic effects.

Fourth, schizophrenia and depression are disorders with an average age of onset around early to mid-adulthood^73,74^. Our measure of lifetime smoking was generated using participants in the UK Biobank aged over 40 years. Therefore, the causal pathway from smoking to schizophrenia and depression risk might initially seem unclear. However, we were not using participants’ smoking behaviour at age 40, but rather retrospective lifetime smoking behaviour from age at initiation. It is plausible that smoking behaviour in earlier life could increase risk of later mental health outcomes and exacerbate symptoms, consequently causing more smokers than non-smokers to seek a diagnosis. We also conducted a sensitivity analysis using smoking initiation as the exposure. Individuals are more likely to initiate smoking prior to the average age at onset of schizophrenia and depression, with 90% of individuals initiating before 18 years of age^75^. With smoking initiation as the exposure, we found consistent evidence that increased likelihood of smoking initiation increased risk of both depression and schizophrenia. There has also been recent debate in the field about the interpretation of time varying exposures in MR and one way to minimise bias is to use average SNP effects on phenotype across time, as we have done here with our lifetime smoking instrument^60^. We recommend that future studies wishing to examine the effects of smoking on an outcome use multiple instruments for smoking behaviour with consistent evidence across multiple instruments providing the strongest evidence of a causal effect.

Fifth, there is a high degree of zero inflation in the distribution of our lifetime smoking index scores with 54% of the sample being never smokers and therefore receiving a score of zero. We decided not to transform the variable given the desire to have interpretable effect sizes for MR and we decided not to exclude never smokers because our instrument is designed to be used in two-sample MR without the need to stratify into smokers and non-smokers. Despite the zero inflation, we see similar effects for lifetime smoking and smoking initiation suggesting that this has not impaired the score. Sixth, the lifetime smoking score was simulated using all-cause mortality and lung cancer as outcomes. The pattern of association between smoking and lung cancer risk compared with risk for schizophrenia and depression is likely to be different. However, increased mortality amongst individuals with schizophrenia and depression is in large part due to smoking related mortality^6^. The effects were modelled on all-cause mortality and lung cancer but no difference to the best fitting value of half-life was observed. We hope that using all-cause mortality as an outcome makes the lifetime smoking instrument broadly applicable to exploring multiple outcomes. Furthermore, the same effects are observed using smoking initiation as the exposure, which does not include the simulated variable.

Finally, there is known selection bias in the UK Biobank sample, with participants being more highly educated, less likely to be a smoker and overall healthier than the general UK population^76^. Of the 9 million individuals contacted, only ~5% consented to take part^76^. Due to the lack of representativeness in the UK Biobank sample, prevalence and incidence rates will not reflect underlying population levels and there is potential for collider bias. If both smoking and risk for schizophrenia and depression reduce the likelihood of participating in the UK Biobank, then this would induce a negative correlation between schizophrenia or depression and smoking. That is the opposite of the effects observed, suggesting our estimates may, if anything, be conservative.

### Conclusion

In conclusion, we have developed a novel genetic instrument of lifetime smoking behaviour that can be used in two-sample MR of summary data without stratifying on smoking status or reducing power. In two-sample MR analysis, there was evidence of an effect between smoking and mental health in both directions; however, the self-medication effects were stronger for depression than schizophrenia. Strong effects of smoking as a risk factor emphasises the detrimental public health consequences of smoking, especially for mental health, and the need to reduce smoking prevalence not only to reduce the burden of physical illness, but also to reduce the burden of mental illness.

## Acknowledgements

We are grateful to the participants of the UK Biobank, the Avon Longitudinal Study of Parents and Children and the individuals who contributed to each of the previous GWAS analyses conducted as well as all the research staff who worked on the data collection.

## Data and code availability

GWAS summary data for lifetime smoking will be made available following publication.

## Author contribution statement

Contributors MRM conceived the study. REW conducted the analysis with statistical assistance from BGS and theoretical assistance from HMS, RBL and MRM. RCR conducted the two-sample MR analysis of lifetime smoking on methylation and performed sensitivity analyses. REW and MRM drafted the initial manuscript. GH assisted with sensitivity analyses. AET provided access to the UK Biobank data. All authors guided the analysis, assisted with interpretation, commented on drafts of the manuscript and approved the final version.

## References

1. World Health Organization. WHO report on the global tobacco epidemic. (World Health Organization, 2011).

2. Coulthard, M., Farrell, M., Singleton, N. & H. Meltzer Tobacco, alcohol and drug use and mental health. Lond. Station. Off. (2002).

3. Lasser, K. et al. Smoking and mental illness: A population-based prevalence study. JAMA 284, 2606–2610 (2000).

4. Lawrence, D., Mitrou, F. & Zubrick, S. R. Smoking and mental illness: results from population surveys in Australia and the United States. BMC Public Health 9, 285 (2009).

5. McClave, A. K., McKnight-Eily, L. R., Davis, S. P. & Dube, S. R. Smoking characteristics of adults with selected lifetime mental illnesses: results from the 2007 National Health Interview Survey. Am. J. Public Health 100, 2464–2472 (2010).

6. Royal College of Physicians. Royal College of Physicians, Royal College of Psychiatrists. Smoking and mental health. (2013).

7. Byers, A. L. et al. Twenty-year depressive trajectories among older women. Arch. Gen. Psychiatry 69, 1073–1079 (2012).

8. Leung, J., Gartner, C., Dobson, A., Lucke, J. & Hall, W. Psychological distress is associated with tobacco smoking and quitting behaviour in the Australian population: evidence from national cross-sectional surveys. Aust. N. Z. J. Psychiatry 45, 170–178 (2011).

9. Tjora, T. et al. The association between smoking and depression from adolescence to adulthood. Addiction 109, 1022–1030 (2014).

10. Adult smoking habits in the UK - Office for National Statistics. Available at: https://www.ons.gov.uk/peoplepopulationandcommunity/healthandsocialcare/healthandlifeexpectancies/bulletins/adultsmokinghabitsingreatbritain/2016#main-points. (Accessed: 9th July 2018)

11. Chang, C.-K. et al. Life Expectancy at Birth for People with Serious Mental Illness and Other Major Disorders from a Secondary Mental Health Care Case Register in London. PLOS ONE 6, e19590 (2011).

12. Khantzian, E. The self-medication hypothesis of substance use disorders: a reconsideration and recent applications. Harv. Rev. Psychiatry 4, 231–244 (1997).

13. Desai, H. D., Seabolt, J. & Jann, M. W. Smoking in patients receiving psychotropic medications. CNS Drugs 15, 469–494 (2001).

14. Levin, E. D., Wilson, W., Rose, J. E. & McEvoy, J. Nicotine–haloperidol interactions and cognitive performance in schizophrenics. Neuropsychopharmacology 15, 429 (1996).

15. Lerman, C. et al. Investigation of mechanisms linking depressed mood to nicotine dependence. Addict. Behav. 21, 9–19 (1996).

16. Lawlor, D. A., Harbord, R. M., Sterne, J. A., Timpson, N. & Davey Smith, G. Mendelian randomization: using genes as instruments for making causal inferences in epidemiology. Stat. Med. 27, 1133–1163 (2008).

17. Ripke, S. et al. Biological insights from 108 schizophrenia-associated genetic loci. Nature 511, 421–427 (2014).

18. Munafò, M. R. et al. Association Between Genetic Variants on Chromosome 15q25 Locus and Objective Measures of Tobacco Exposure. JNCI J. Natl. Cancer Inst. 104, 740–748 (2012).

19. Thorgeirsson, T. E. et al. A Variant Associated with Nicotine Dependence, Lung Cancer and Peripheral Arterial Disease. Nature 452, 638–642 (2008).

20. Tobacco Consortium. Genome-wide meta-analyses identify multiple loci associated with smoking behavior. Nat. Genet. 42, 441–447 (2010).

21. Ware, J. J., van den Bree, M. B. & Munafò, M. R. Association of the CHRNA5-A3-B4 gene cluster with heaviness of smoking: a meta-analysis. Nicotine Tob. Res. 13, 1167–1175 (2011).

22. Gage, S. H. & Munafò, M. R. Rethinking the association between smoking and schizophrenia. Lancet Psychiatry 2, 118 (2015).

23. Hartz, S. M. et al. Genetic correlation between smoking behaviors and schizophrenia. Schizophr. Res. 194, 86–90 (2018).

24. Reginsson, G. W. et al. Polygenic risk scores for schizophrenia and bipolar disorder associate with addiction. Addict. Biol. 23, 485–492 (2018).

25. Kendler, K. S., Lönn, S. L., Sundquist, J. & Sundquist, K. Smoking and schizophrenia in population cohorts of Swedish women and men: a prospective co-relative control study. Am. J. Psychiatry 172, 1092–1100 (2015).

26. Kendler, K. S. et al. Smoking and major depression: a causal analysis. Arch. Gen. Psychiatry 50, 36–43 (1993).

27. Gurillo, P., Jauhar, S., Murray, R. M. & MacCabe, J. H. Does tobacco use cause psychosis? Systematic review and meta-analysis. Lancet Psychiatry 2, 718–725 (2015).

28. Scott, J. G. et al. Evidence of a Causal Relationship Between Smoking Tobacco and Schizophrenia Spectrum Disorders. Front. Psychiatry 9, (2018).

29. Taylor, G. et al. Change in mental health after smoking cessation: systematic review and meta-analysis. BMJ 348, g1151 (2014).

30. Munafò, M. R. & Smith, G. D. Robust research needs many lines of evidence. Nature (2018). doi:10.1038/d41586-018-01023-.

