## Supplementary Materials for "Causal effects of lifetime smoking on risk for depression and schizophrenia: Evidence from a Mendelian randomisation study"

| Supplementary Note  Manuscript references continued | Page 2  Page 4 |
| --- | --- |
| **Figures**  Figure S1. Quantile-Quantile Plot of lifetime smoking index | Page 9 |
| Figure S2. Scatter plot of IVW and sensitivity analyses of lifetime smoking on coronary artery disease | Page 10 |
| Figure S3. Scatter plot of IVW and sensitivity analyses of lifetime smoking on lung cancer | Page 11 |
| Figure S4. Single SNP analysis of lifetime smoking on coronary artery disease | Page 12 |
| Figure S5. Single SNP analysis of lifetime smoking on lung cancer | Page 13 |
| Figure S6. Leave-one-out analysis of lifetime smoking on coronary artery disease | Page 14 |
| Figure S7. Leave-one-out SNP analysis of lifetime smoking on lung cancer | Page 15 |
| Figure S8. Scatter plot of IVW and sensitivity analyses of lifetime smoking on schizophrenia | Page 16 |
| Figure S9. Scatter plot of IVW and sensitivity analyses of lifetime smoking on major depressive disorder | Page 17 |
| Figure S10. Scatter plot of IVW and sensitivity analyses of schizophrenia on lifetime smoking | Page 18 |
| Figure S11. Scatter plot of IVW and sensitivity analyses of depression on lifetime smoking | Page 19 |
| Figure S12. Single SNP analysis of lifetime smoking on schizophrenia | Page 20 |
| Figure S13. Single SNP analysis of lifetime smoking on depression | Page 21 |
| Figure S14. Single SNP analysis of schizophrenia on lifetime smoking | Page 22 |
| Figure S15. Single SNP analysis of depression on lifetime smoking | Page 23 |
| Figure S16. Leave-one-out analysis of lifetime smoking on schizophrenia | Page 24 |
| Figure S17. Leave-one-out analysis of lifetime smoking on depression | Page 25 |
| Figure S18. Leave-one-out analysis of schizophrenia on lifetime smoking | Page 26 |
| Figure S19. Leave-one-out analysis of depression on lifetime smoking | Page 27 |
| **Tables** |  |
| Table S1. SNPs associated with lifetime smoking index at the genome-wide level of significance | Page 28 |
| Table S2. Tests of the unweighted and weighted regression dilution I^2^_GX_ for the lifetime smoking exposure  Table S3. Tests of heterogeneity | Page 32  Page 33 |
| Table S4. Test of directional pleiotropy using the MR Egger Intercept  Table S5. Mendelian randomisation results following Steiger filtering. | Page 34  Page 36 |
| Table S6. Bi-directional two-sample Mendelian randomisation analyses of the effect of lifetime smoking on depression (2013).  Table S7. Sensitivity analysis without genotyping chip used as a covariate.  Table S8. Sensitivity analysis with smoking initiation GWAS from GSCAN.  Table S9. Bi-directional two-sample Mendelian randomisation analyses of the effect of lifetime smoking on schizophrenia (2018).  Table S10. Comparison of MR Sensitivity methods | Page 37  Page 38  Page 39  Page 40  Page 41  Page 43 |
| **References** |  |

**Supplementary Note**

**Construction of the lifetime smoking index.**

We simulated values of two constants: half-life ($\tau$), and lag time ($\delta$). Together these constants capture the non-linear risk of smoking on health. Half-life captures the exponentially decreasing effect of smoking at a given time on health outcomes. Lag time accounts for the observation that smokers are more at risk of certain diseases (e.g., lung cancer) immediately after stopping smoking than current smokers^1^. This is likely due to the prodromal consequences of disease being felt by the individual.

Simulations were run for possible values of τ (between 2 and 50 varying in increments of 1) and δ (between 0 and 5, varying in increments of 0.1) to find the best fitting model to explain the effects of lifetime smoking on lung cancer and all-cause mortality. Akaike information criterion (AIC) was used to select the best fitting model. The best fitting value for half-life was 18 for both lung cancer and all-cause mortality. We did not see any improvement of fit for changes in the value of lag time, therefore this was set to 0 as has been done previously ^2^. This removes the effects of lag time from the model. These values were used to fit the final model which is:

tsc* = max(tsc - δ, 0)

dur* = max(dur + tsc - δ, 0) – tsc*

lifetime smoking = (1 – 0.5^dur*/τ^) (0.5^tsc*/τ^) ln(int+1)

…where τ = half-life, δ = lag time, int = cigarettes per day, tss = time started smoking, tsc = time since cessation, dur = duration of smoking (either age-tss for current smokers or [age-tsc]-tss for former smokers). So, in our case, where δ = 0, tsc* = tsc and consequently, our lifetime smoking calculation can be simplified to:

lifetime smoking = (1 – 0.5^dur/τ^) (0.5^tsc/τ^) ln(int+1)

Individuals who have never smoked and have no smoking exposure will have a value of 0. In our sample, values for smokers ranged from 0.007 (an individual who smoked 1 cigarette per day for 1 year) to 4.169 (an individual who currently smokes 140 a day and started smoking at age 11 years of age). Values of lifetime smoking were treated as continuous in subsequent analysis.

**Genotyping and exclusion procedure.**

UK Biobank participants provided blood samples at initial assessment centre. Genotyping was performed using the Affymetrix UK BiLEVE Axiom array for 49,979 participants and using the Affymetrix UK Biobank Axiom® array for 438,398 participants. The two arrays share 95% coverage, but chip is adjusted for in all analyses because the UK BiLEVE sample is over represented for smokers. Imputation and initial quality control steps were performed by the Wellcome Trust Centre for Human Genetics resulting in over 90 million single nucleotide polymorphisms (SNPs) and indels^3^.

Individuals were excluded if there were sex-mismatches between reported and chromosomal sex or aneuploidy (N=814). Individuals were restricted to European ancestry based on the first four principal components of population structure and related individuals were removed following MRC Integrative Epidemiology Unit filtering steps^4^. After excluding individuals who had withdrawn consent, 463,033 of the participants remained^4^. We restricted our analysis to autosomes only and used filtering thresholds for SNPs of minor allele frequency (MAF) >0.01 and info score (measure of imputation uncertainty) >0.8.

**ALSPAC sample description and measures.**

To test prediction in an independent sample, we used 2,712 mothers from Avon Longitudinal Study of Parents and Children (ALSPAC). ALSPAC is a longitudinal birth cohort, which recruited 14,541 pregnant women between 1991 and 1992 with detailed descriptions reported elsewhere^5,6^. Self-reported measures of smoking status, age at initiation, age at cessation and cigarettes per day were collected from these women when their offspring were 18 years of age (mean mothers age = 48 years, SD = 4.3).

**Outcome GWAS samples for instrument validation.**

For lung cancer, we used the summary data from the ILCCO consortium GWAS, which comprises 11,348 cases and 15,861 controls of European ancestry^7^. Lung cancer cases were classified by tumour type using either the International Classification of Diseases for Oncology (ICD-O) or the World Health Organisation (WHO) coding. Tumours with overlapping histologies were classified as mixed. Most samples in the GWAS meta-analysis only included adenocarcinomas (AD) or squamous carcinomas (SQ) but classification was different for each contributing cohort. The GWAS was run with a binary outcome, case control status for any primary lung cancer tumour. For more details see Wang et al. (2014)^7^.

For CAD, we used the GWAS summary data from CARDIoGRAMplusC4D which comprises 60,801 cases and 123,504 controls of mixed ancestry^8^. Case status was defined as any CAD diagnosis including myocardial infarction, acute coronary syndrome, chronic stable angina or coronary stenosis of >50%. This GWAS was conducted with a binary outcome of CAD cases compared with controls.

We conducted a GWAS of AHRR (cg05575921) methylation in the Accessible Resource for Integrated Epigenomic Studies (ARIES) subset of ALSPAC^9^. Genome-wide DNA methylation profiling in ARIES was performed using the Illumina Infinium HumanMethylation450 BeadChip (450K) array for ~1000 mother-offspring pairs^9^. For this analysis, we used methylation data derived from whole blood which was collected from mothers (846 smokers and non-smokers) when the index offspring were around 18 years of age. Methylation data were normalised in R with the wateRmelon package^10^ using the Touleimat and Tost^11^algorithm to reduce the non-biological differences between probes. As was done previously^12^, AHRR methylation (cg05575921) was rank-normalised to remove outliers and regressed on the following covariates: age, the top ten ancestry principal components, bisulphite conversion batch and estimated white blood cell counts (using an algorithm based on differential methylation between cell types^13^). Residuals were then taken forward and SNP association effects were obtained in PLINK1.07 using exact linear regression. Full details of the GWAS methods used for AHRR locus methylation are described elsewhere^9^.

**Manuscript references continued**

31.	Davey Smith, G. & Ebrahim, S. ‘Mendelian randomization’: can genetic epidemiology contribute to understanding environmental determinants of disease? *Int. J. Epidemiol.* **32**, 1–22 (2003).

32.	Davey Smith, G. & Hemani, G. Mendelian randomization: genetic anchors for causal inference in epidemiological studies. *Hum. Mol. Genet.* **23**, R89–R98 (2014).

33.	Bjørngaard, J. H. *et al.* The causal role of smoking in anxiety and depression: a Mendelian randomization analysis of the HUNT study. *Psychol. Med.* **43**, 711–719 (2013).

34.	Taylor, A. E. *et al.* Investigating the possible causal association of smoking with depression and anxiety using Mendelian randomisation meta-analysis: the CARTA consortium. *BMJ Open* **4**, e006141 (2014).

35.	Wium-Andersen, M. K., Ørsted, D. D. & Nordestgaard, B. G. Tobacco smoking is causally associated with antipsychotic medication use and schizophrenia, but not with antidepressant medication use or depression. *Int. J. Epidemiol.* **44**, 566–577 (2015).

36.	Gage, S. H. *et al.* Investigating causality in associations between smoking initiation and schizophrenia using Mendelian randomization. *Sci. Rep.* **7**, 40653 (2017).

37.	Leffondré, K., Abrahamowicz, M., Xiao, Y. & Siemiatycki, J. Modelling smoking history using a comprehensive smoking index: application to lung cancer. *Stat. Med.* **25**, 4132–4146 (2006).

38.	Elsworth, B. L. *et al.* MRC IEU UK Biobank GWAS pipeline version 1. *data.bris* (2017). doi:10.5523/bris.2fahpksont1zi26xosyamqo8rr

39.	Loh, P.-R. *et al.* Efficient Bayesian mixed-model analysis increases association power in large cohorts. *Nat. Genet.* **47**, 284–290 (2015).

40.	Hemani, G. *et al.* The MR-Base platform supports systematic causal inference across the human phenome. *eLife* **7**, e34408 (2018).

41.	Purcell, S. *et al.* PLINK: a tool set for whole-genome association and population-based linkage analyses. *Am. J. Hum. Genet.* **81**, 559–575 (2007).

42.	Zeilinger, S. *et al.* Tobacco Smoking Leads to Extensive Genome-Wide Changes in DNA Methylation. *PLOS ONE* **8**, e63812 (2013).

43.	R. Core Team. *R: A language and environment for statistical computing. R Foundation for Statistical Computing, Vienna, Austria. 2013*. (ISBN 3-900051-07-0, 2014).

44.	Bowden, J., Davey Smith, G. & Burgess, S. Mendelian randomization with invalid instruments: effect estimation and bias detection through Egger regression. *Int. J. Epidemiol.* **44**, 512–525 (2015).

45.	Bowden, J., Davey Smith, G., Haycock, P. C. & Burgess, S. Consistent estimation in Mendelian randomization with some invalid instruments using a weighted median estimator. *Genet. Epidemiol.* **40**, 304–314 (2016).

46.	Hartwig, F. P., Smith, G. D. & Bowden, J. Robust inference in two-sample Mendelian randomisation via the zero modal pleiotropy assumption. *bioRxiv* 126102 (2017).

47.	Zhao, Q., Wang, J., Hemani, G., Bowden, J. & Small, D. S. Statistical inference in two-sample summary-data Mendelian randomization using robust adjusted profile score. *arXiv* (2018).

48.	Lawlor, D. A., Tilling, K. & Davey Smith, G. Triangulation in aetiological epidemiology. *Int. J. Epidemiol.* **45**, 1866–1886 (2016).

49.	Pardiñas, A. F. *et al.* Common schizophrenia alleles are enriched in mutation-intolerant genes and in regions under strong background selection. *Nat. Genet.* **50**, 381–389 (2018).

50.	Wray, N. R., Sullivan, P. F. & others. Genome-wide association analyses identify 44 risk variants and refine the genetic architecture of major depression. *Nat. Genet.* **50**, 668–681 (2018).

51.	Liu, M. Association studies of up to 1.2 million individuals yield new insights in the genetic etiology of tobacco and alcohol use. *Nat. Genet.* (In Press).

52.	Hemani, G., Tilling, K. & Davey Smith, G. Orienting the causal relationship between imprecisely measured traits using GWAS summary data. *PLOS Genet.* **13**, e1007081 (2017).

53.	Furberg, H. *et al.* Genome-wide meta-analyses identify multiple loci associated with smoking behavior. *Nat. Genet.* **42**, 441 (2010).

54.	Bowden, J. *et al.* Assessing the suitability of summary data for two-sample Mendelian randomization analyses using MR-Egger regression: the role of the I 2 statistic. *Int. J. Epidemiol.* **45**, 1961–1974 (2016).

55.	Burgess, S. *et al.* Using published data in Mendelian randomization: a blueprint for efficient identification of causal risk factors. *Eur. J. Epidemiol.* **30**, 543–552 (2015).

56.	Ripke, S. *et al.* A mega-analysis of genome-wide association studies for major depressive disorder. *Mol. Psychiatry* **18**, 497–511 (2013).

57.	Jones, H. J. *et al.* Association of Combined Patterns of Tobacco and Cannabis Use in Adolescence With Psychotic Experiences. *JAMA Psychiatry* **75**, 240–246 (2018).

58.	Zammit, S. *et al.* Investigating the association between cigarette smoking and schizophrenia in a cohort study. *Am. J. Psychiatry* **160**, 2216–2221 (2003).

59.	Cook, B. L. *et al.* Trends in Smoking Among Adults With Mental Illness and Association Between Mental Health Treatment and Smoking Cessation. *JAMA* **311**, 172–182 (2014).

60.	Labrecque, J. A. & Swanson, S. A. Interpretation and Potential Biases of Mendelian Randomization Estimates With Time-Varying Exposures. *Am. J. Epidemiol.* doi:10.1093/aje/kwy204

61.	Jarvik, M. E. *et al.* Nicotine blood levels and subjective craving for cigarettes. *Pharmacol. Biochem. Behav.* **66**, 553–558 (2000).

62.	Parrott, A. C. Does cigarette smoking cause stress? *Am. Psychol.* **54**, 817 (1999).

63.	Parrott, A. C. Cigarette-derived nicotine is not a medicine. *World J. Biol. Psychiatry* **4**, 49–55 (2003).

64.	Hughes, J. R. Effects of abstinence from tobacco: valid symptoms and time course. *Nicotine Tob. Res.* **9**, 315–327 (2007).

65.	Benowitz, N. L. Nicotine Addiction. *N. Engl. J. Med.* **362**, 2295–2303 (2010).

66.	Howes, O., McCutcheon, R. & Stone, J. Glutamate and dopamine in schizophrenia: An update for the 21st century. *J. Psychopharmacol. (Oxf.)* **29**, 97–115 (2015).

67.	Jakubovski, E., Varigonda, A. L., Freemantle, N., Taylor, M. J. & Bloch, M. H. Systematic Review and Meta-Analysis: Dose-Response Relationship of Selective Serotonin Reuptake Inhibitors in Major Depressive Disorder. *Am. J. Psychiatry* **173**, 174–183 (2015).

68.	Hindocha, C. *et al.* Associations between cigarette smoking and cannabis dependence: a longitudinal study of young cannabis users in the United Kingdom. *Drug Alcohol Depend.* **148**, 165–171 (2015).

69.	Moore, T. H. *et al.* Cannabis use and risk of psychotic or affective mental health outcomes: a systematic review. *The Lancet* **370**, 319–328 (2007).

70.	United Nations Office on Drugs and Crime (UNODC). *World Drug Report 2011.* (2011).

71.	Haworth, S. *et al.* Common genetic variants and health outcomes appear geographically structured in the UK Biobank sample: Old concerns returning and their implications. *bioRxiv* 294876 (2018). doi:10.1101/294876

72.	Cai, N., Kendler, K. & Flint, J. Minimal phenotyping yields GWAS hits of low specificity for major depression. *bioRxiv* 440735 (2018).

73.	WHO | Schizophrenia. *WHO* Available at: https://www.who.int/mental_health/management/schizophrenia/en/. (Accessed: 19th December 2018)

74.	Measuring national well-being: domains and measures - Office for National Statistics. Available at: https://www.ons.gov.uk/peoplepopulationandcommunity/wellbeing/datasets/measuringnationalwellbeingdomainsandmeasures. (Accessed: 19th December 2018)

75.	Centers for Disease Control and Prevention. Smoking and Tobacco Use; Tobacco-Related Disparities; Tobacco Use Among Adults with Mental Illness and Substance Use Disorders. *Smoking and Tobacco Use* (2018). Available at: http://www.cdc.gov/tobacco/basic_information/health_disparities/mental-illness-substance-use/. (Accessed: 24th April 2018)

76.	Munafo, M. R., Tilling, K., Taylor, A. E., Evans, D. M. & Smith, G. D. Collider Scope: When selection bias can substantially influence observed associations. *bioRxiv* 079707 (2017).

**Figure 1. Quantile-quantile plot of SNP associations with lifetime smoking index.**

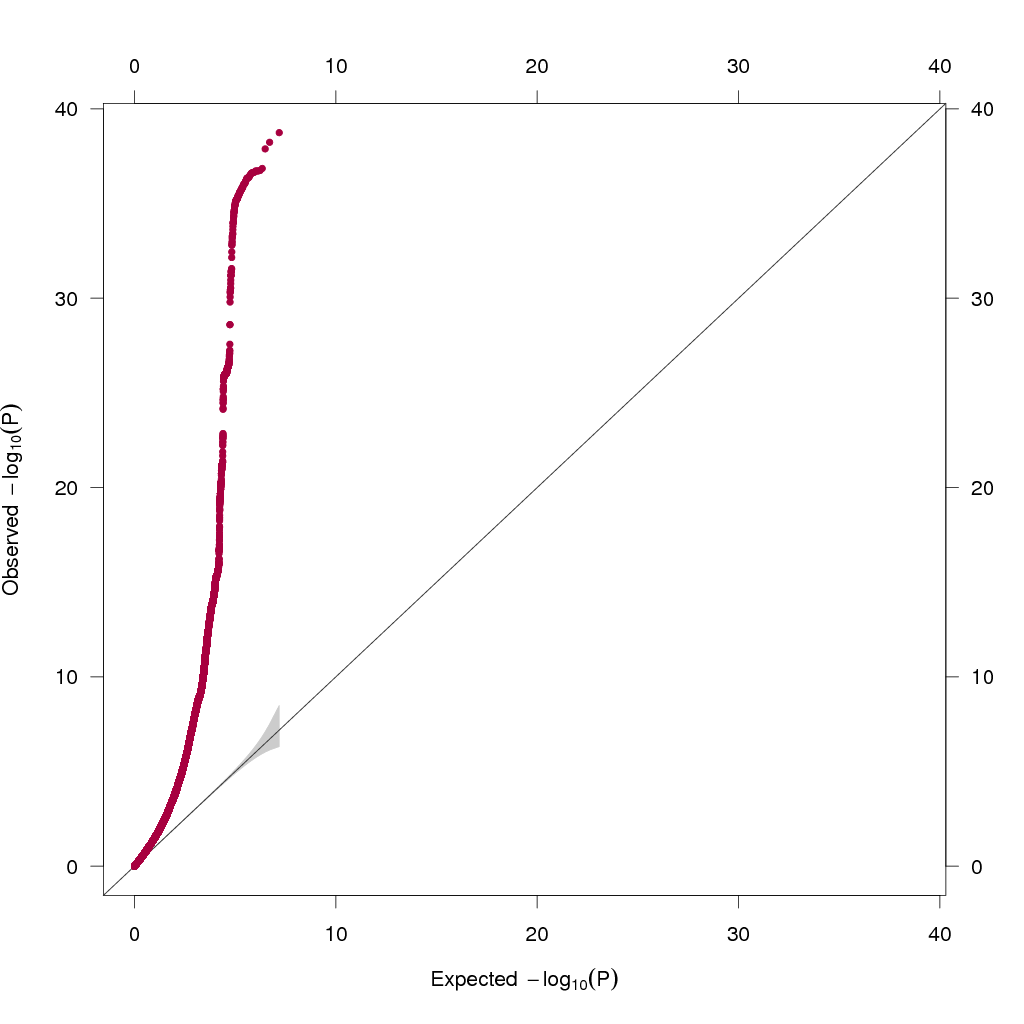

The x-axis indicates the -log_10_ p-value expected under the null hypothesis and the y-axis represents the observed -log_10_ p-value.

Testing inflation by population structure

The QQ plot suggests evidence of inflation due to population stratification. To check this, we calculated the LD score intercept (1.051, SE = 0.0094) and mean chi-square statistic (1.907) which give an attenuation ratio of 0.056, minimal evidence of inflation. The pattern seen here is therefore likely due to the large sample size giving power to detect high numbers of associations.

**Figure S2. Scatter plot of IVW and sensitivity analyses of lifetime smoking on coronary artery disease.**

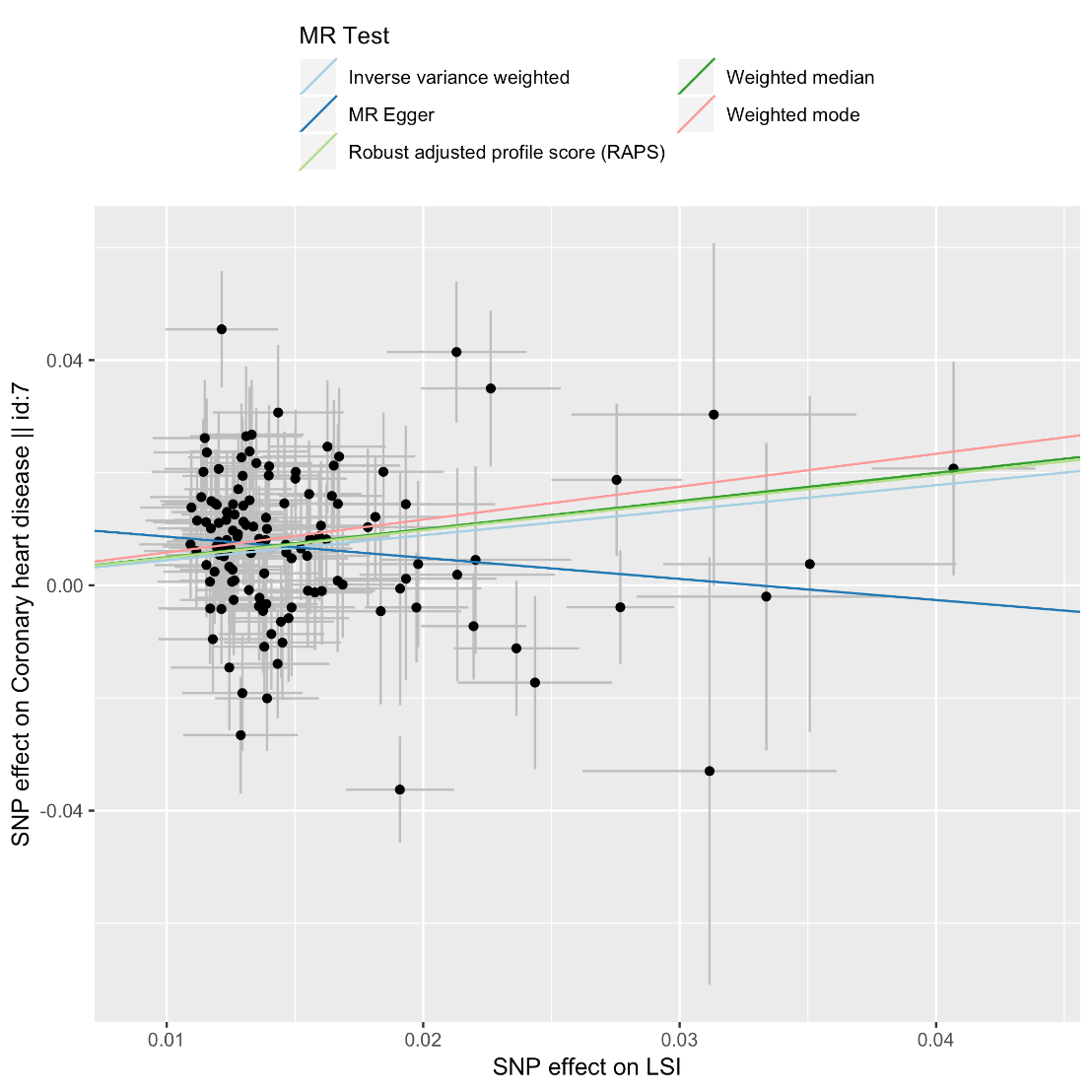

**Figure S3. Scatter plot of IVW and sensitivity analyses of lifetime smoking on lung cancer.**

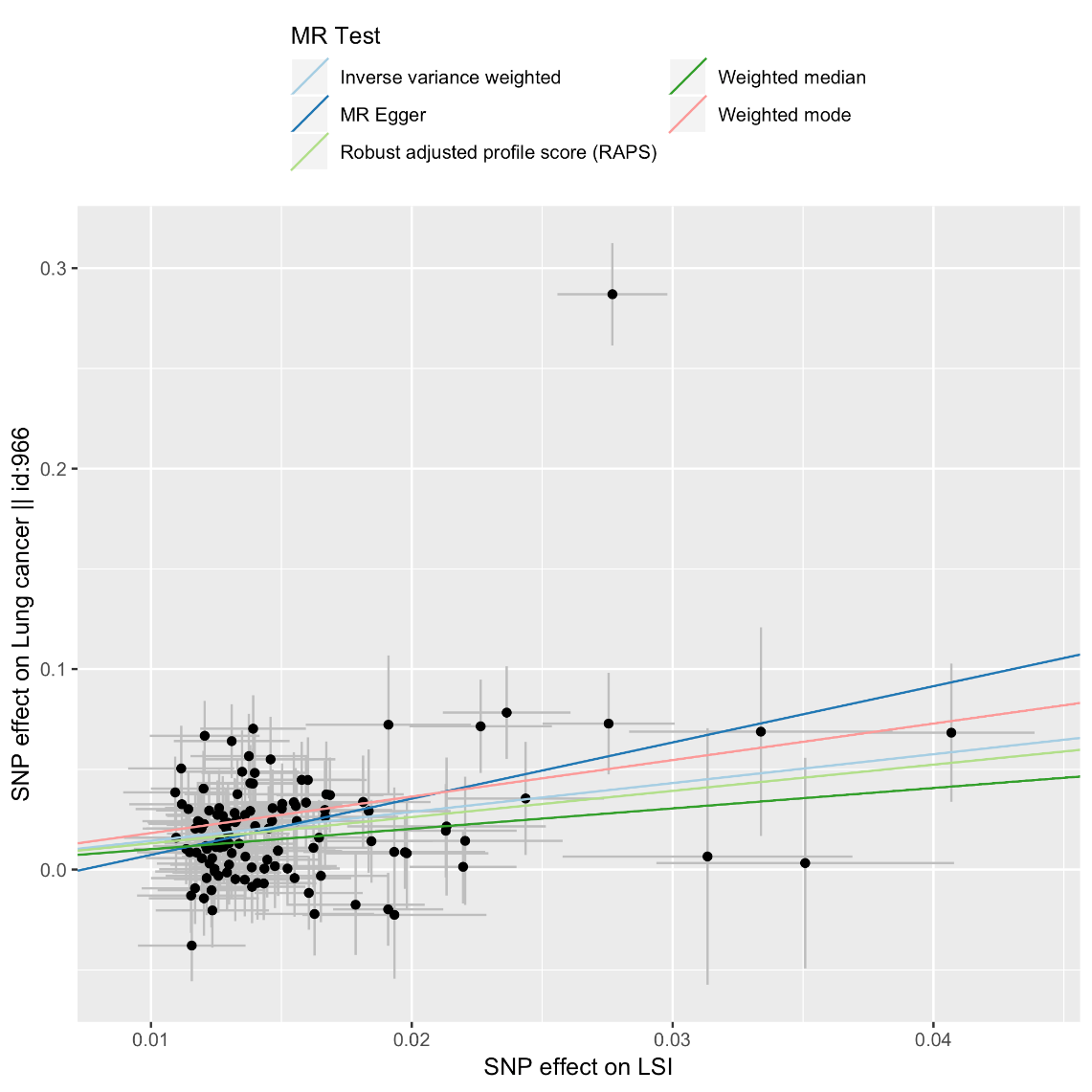

Figure S3 shows evidence of an outlier, however this plot does not account for SE in the SNP effect. Therefore, we followed up this outlier using radial MR.

Radial MR

A radial MR analysis of lifetime smoking on lung cancer using first order weights and an alpha level of 0.01 identified 5 outliers for the IVW method (in order of how large an outlier they are):

**IVW Outliers**

1. rs10918701

2. rs12244388

3. rs329120

4. rs6562474

5. rs8042849

Outliers are usually removed in an incremental fashion, beginning with the largest. However, we can see from our leave-one-out analysis (see Figure S7) that the two largest outliers for both methods (rs10918701 and rs12244388 ) do not affect the estimate once removed.

**Figure S4. Single SNP effects of lifetime smoking on coronary artery disease.**

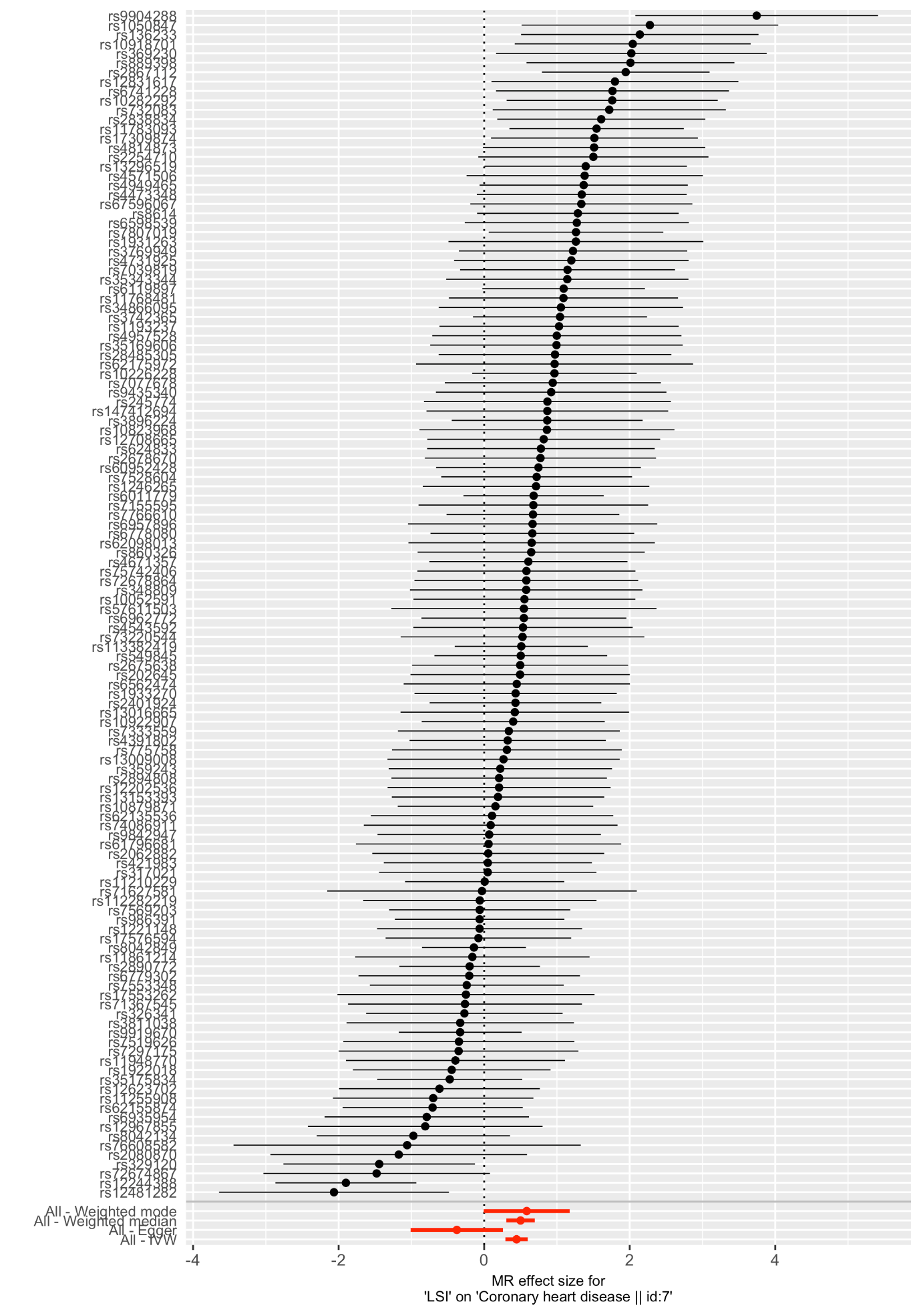

**Figure S5. Single SNP effects of lifetime smoking on lung cancer.**

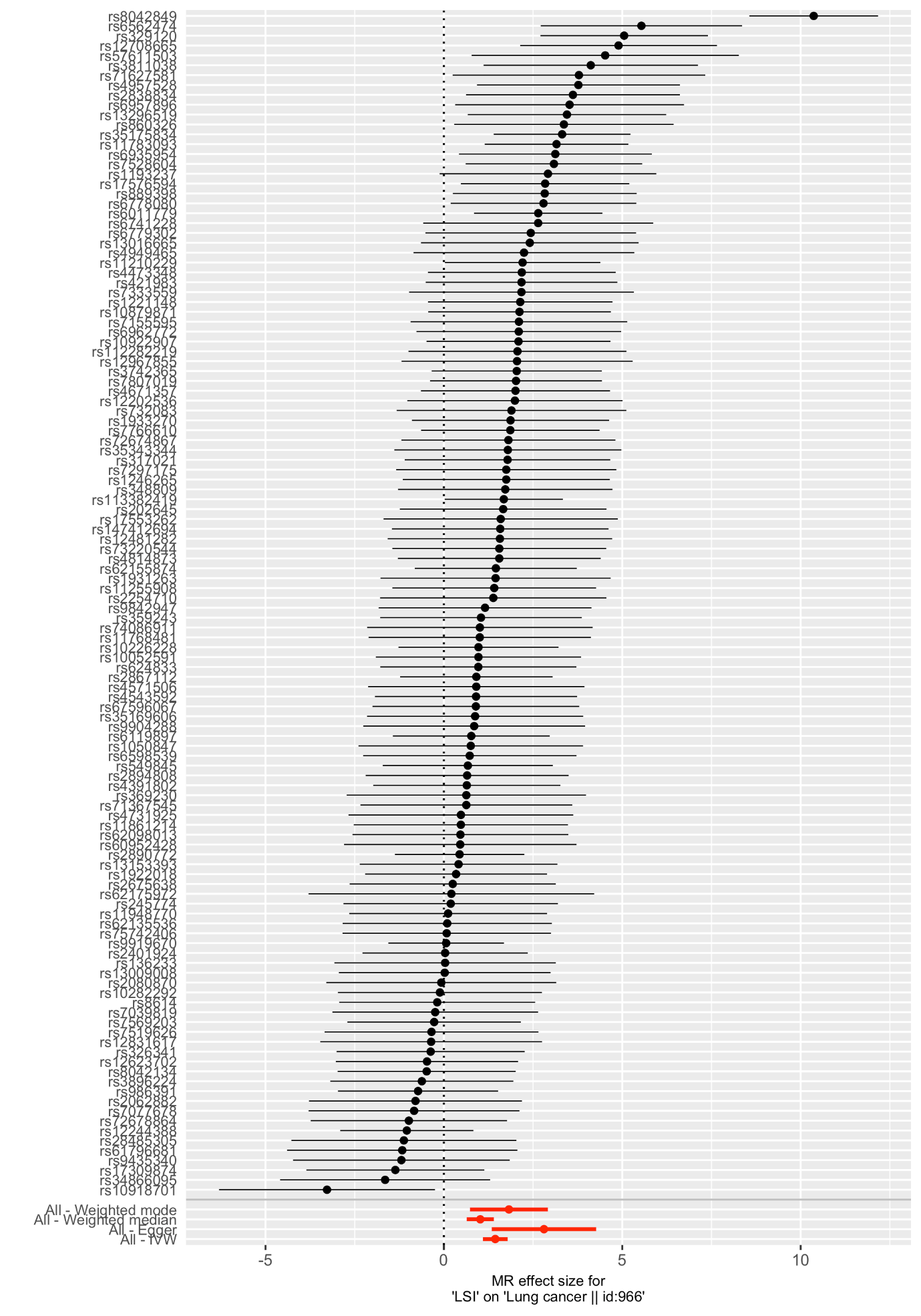

The SNP with the largest effect is rs8042849, which is an intron variant of the *HYKK* gene, has previously been associated with nicotine dependence and forced expiratory volume (FEV).

**Figure S6. Leave-one-out effects of lifetime smoking on coronary artery disease using the inverse-variance weighted method.**

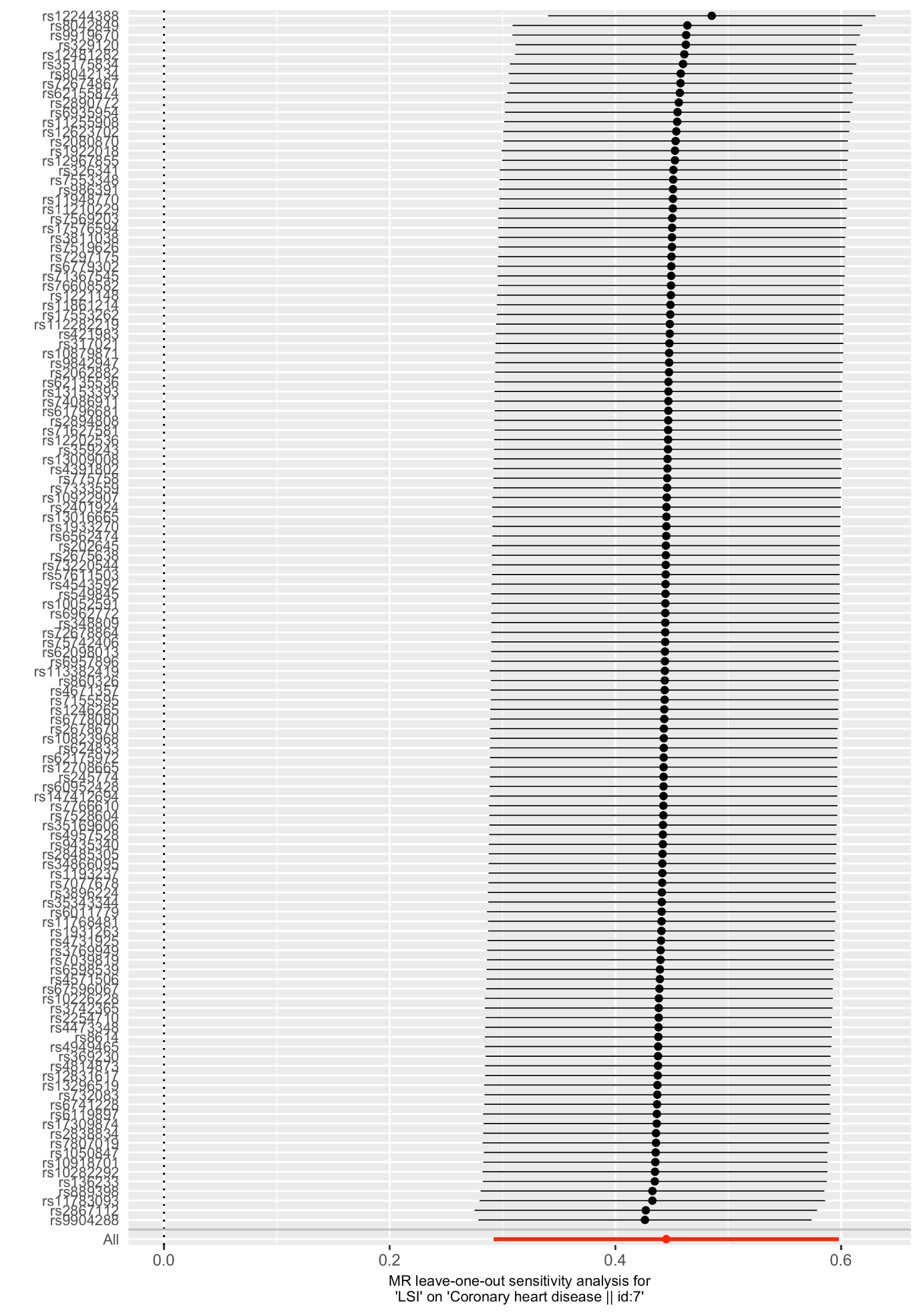

**Figure S7. Leave-one-out effects of lifetime smoking on lung cancer using the inverse-variance weighted method.**

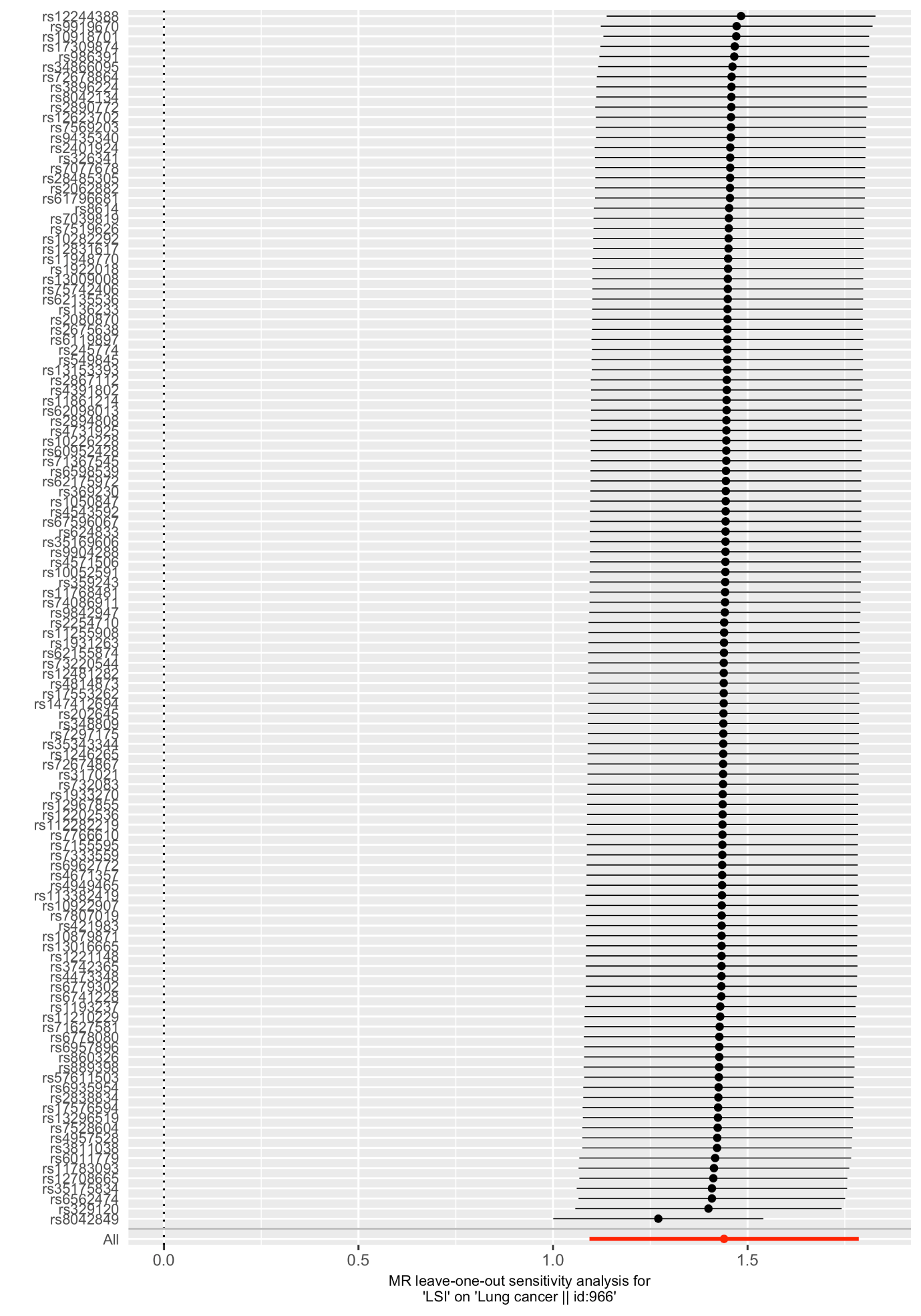

**Figure S8. Scatter plot of IVW and sensitivity analyses of lifetime smoking on schizophrenia.**

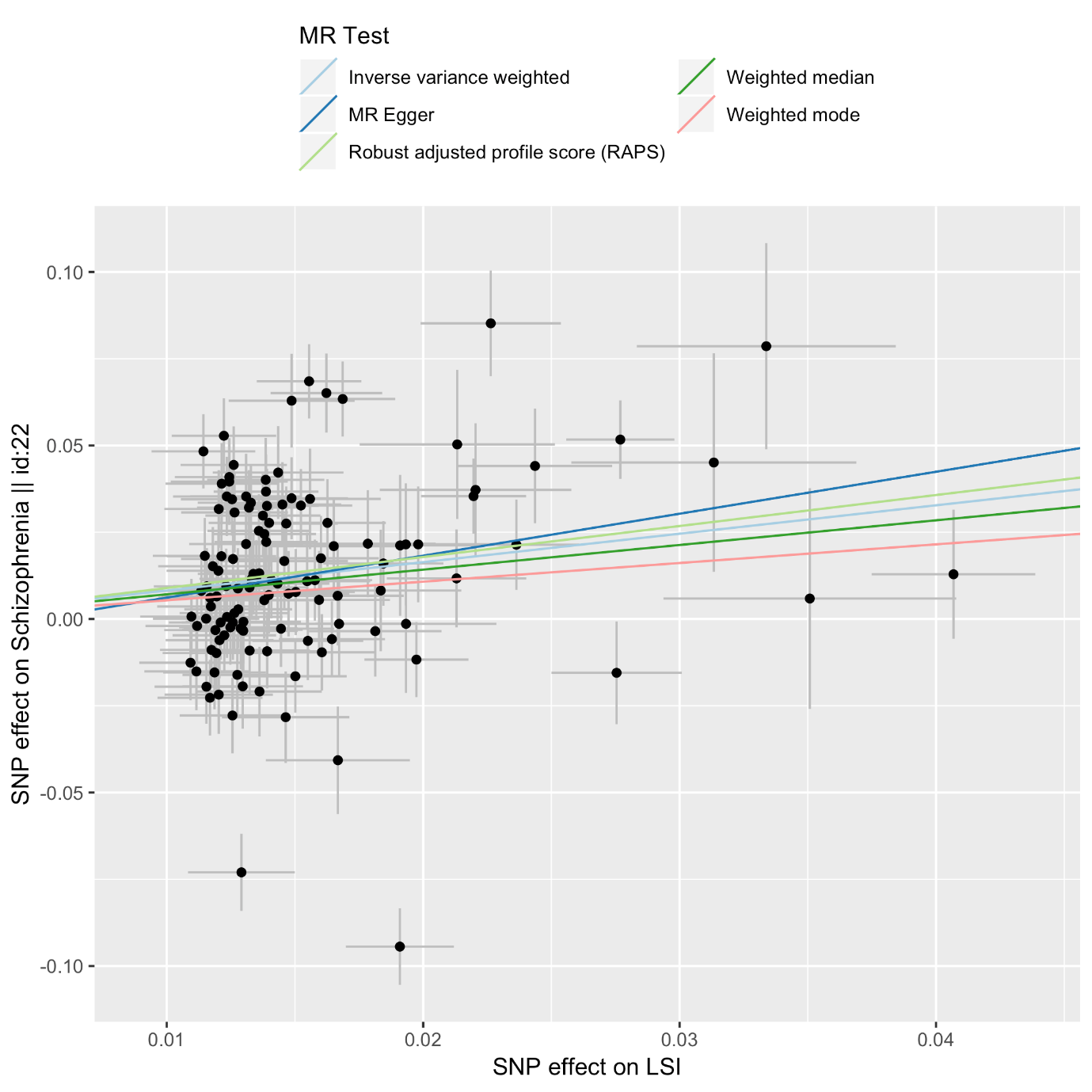

**Figure S9. Scatter plot of IVW and sensitivity analyses of lifetime smoking on major depressive disorder.**

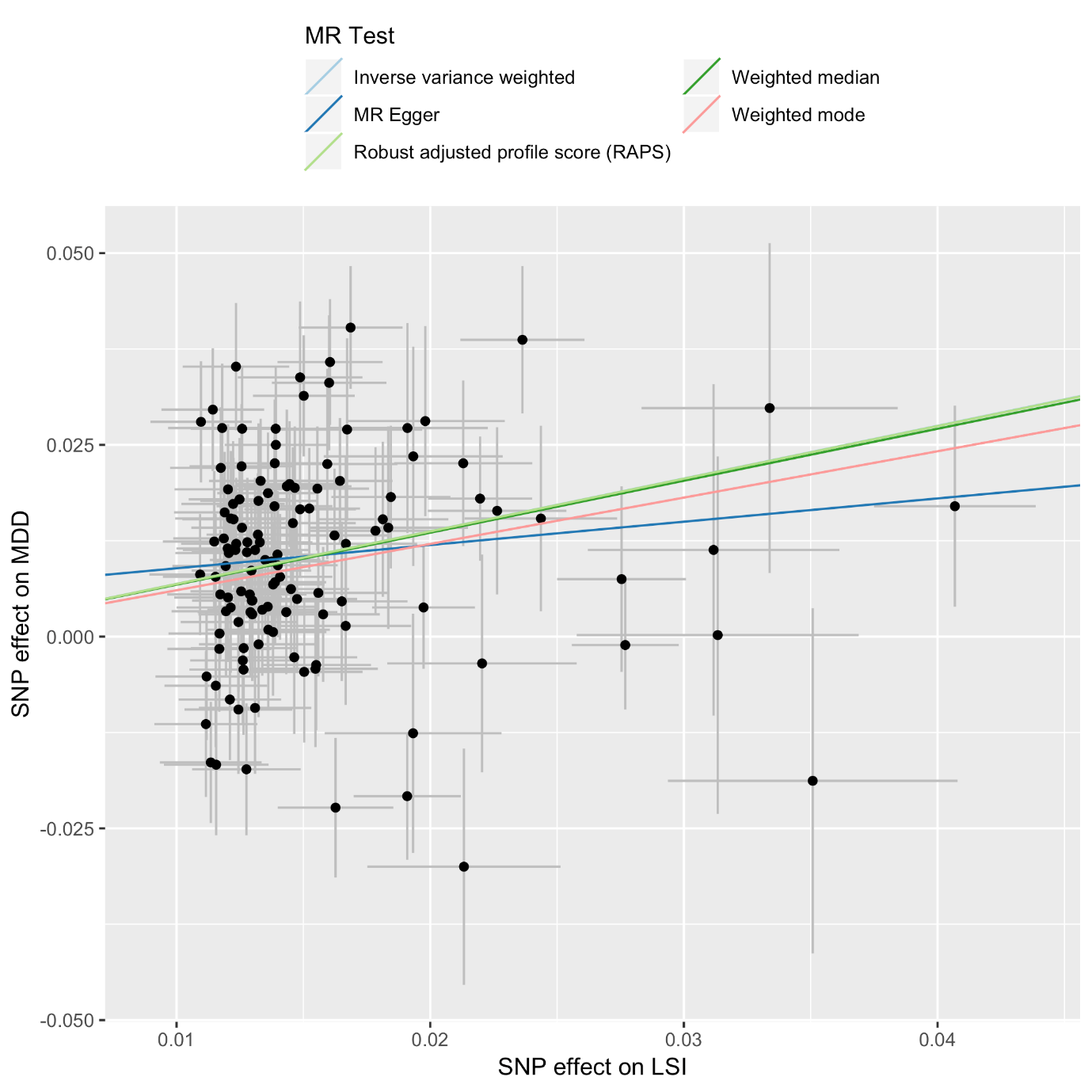

**Figure S10. Scatter plot of IVW and sensitivity analyses of schizophrenia on lifetime smoking.**

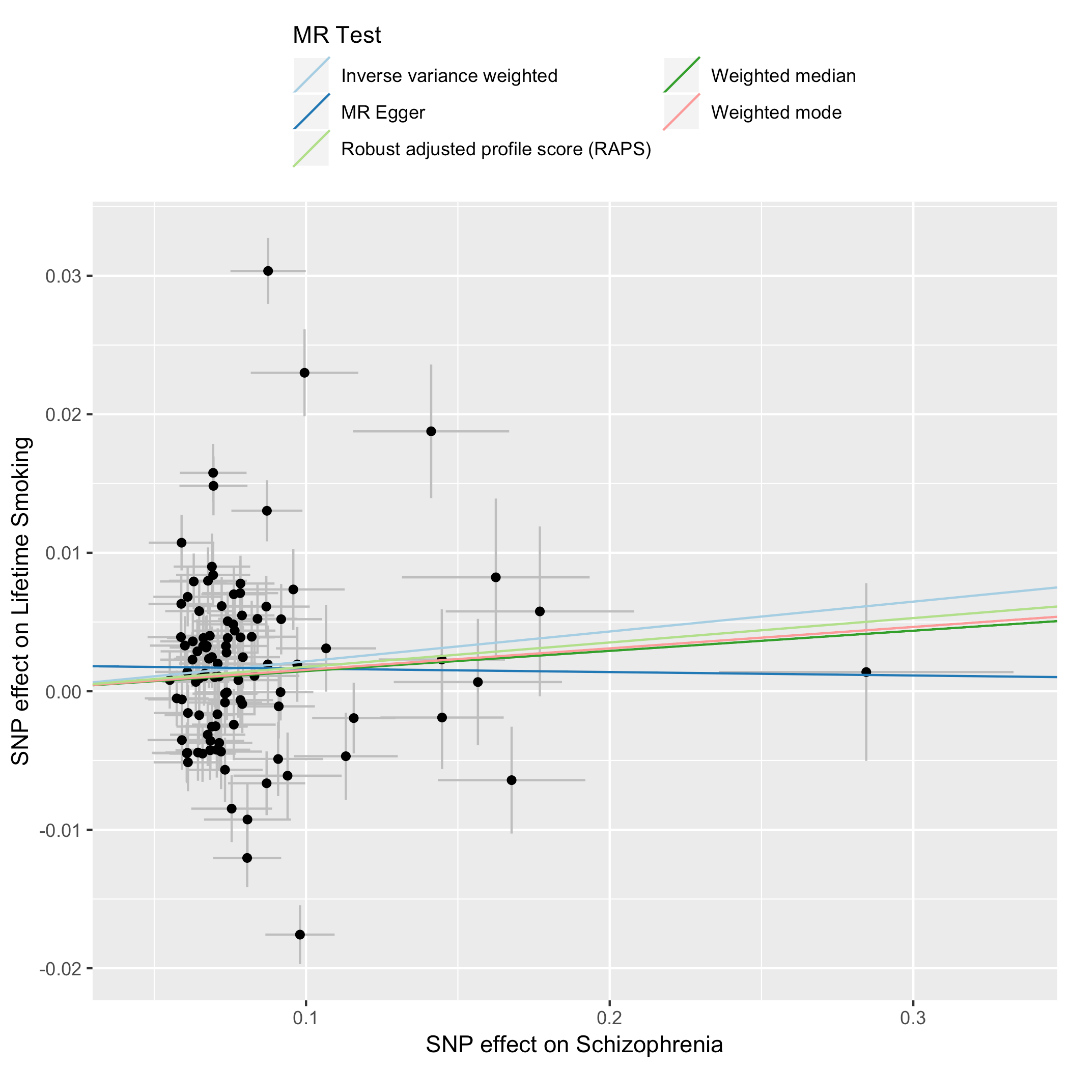

**Figure S11. Scatter plot of IVW and sensitivity analyses of depression on lifetime smoking.**

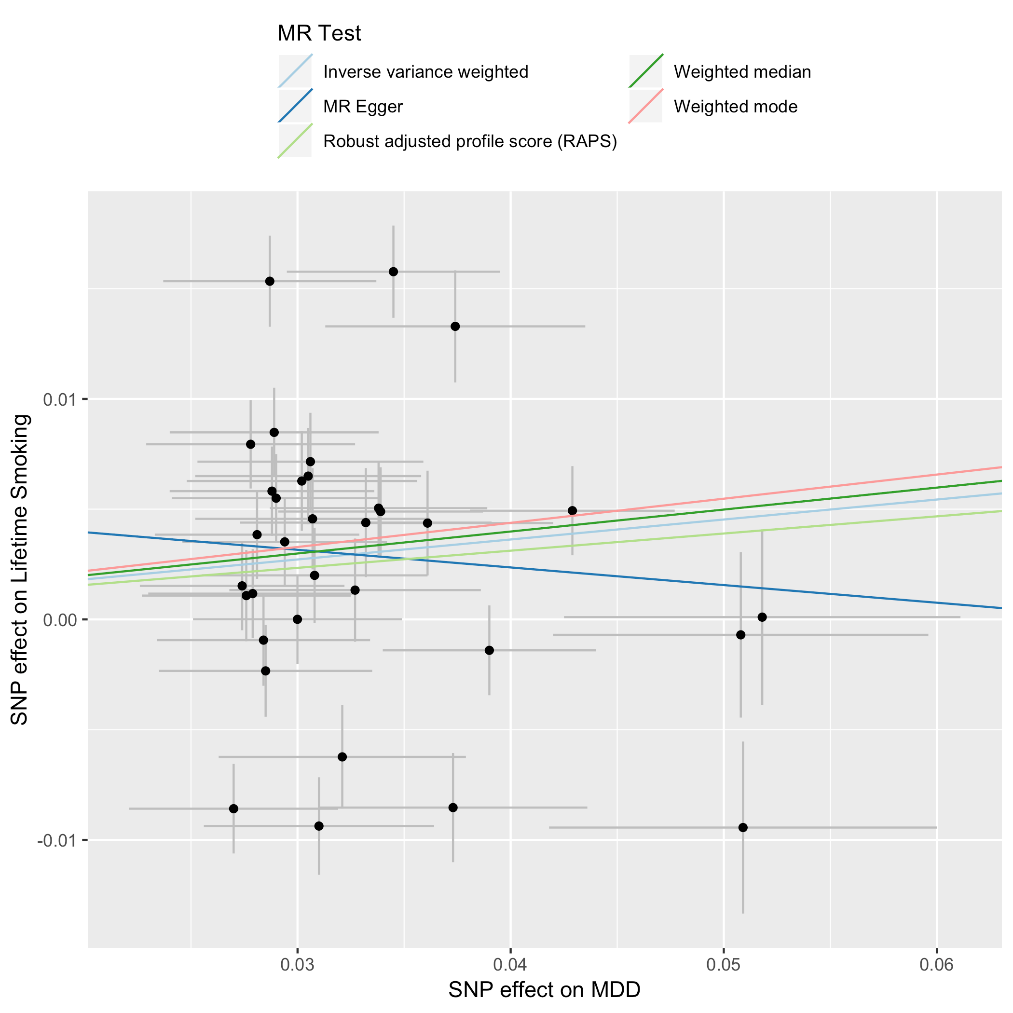

**Figure S12. Single SNP analysis of lifetime smoking on schizophrenia.**

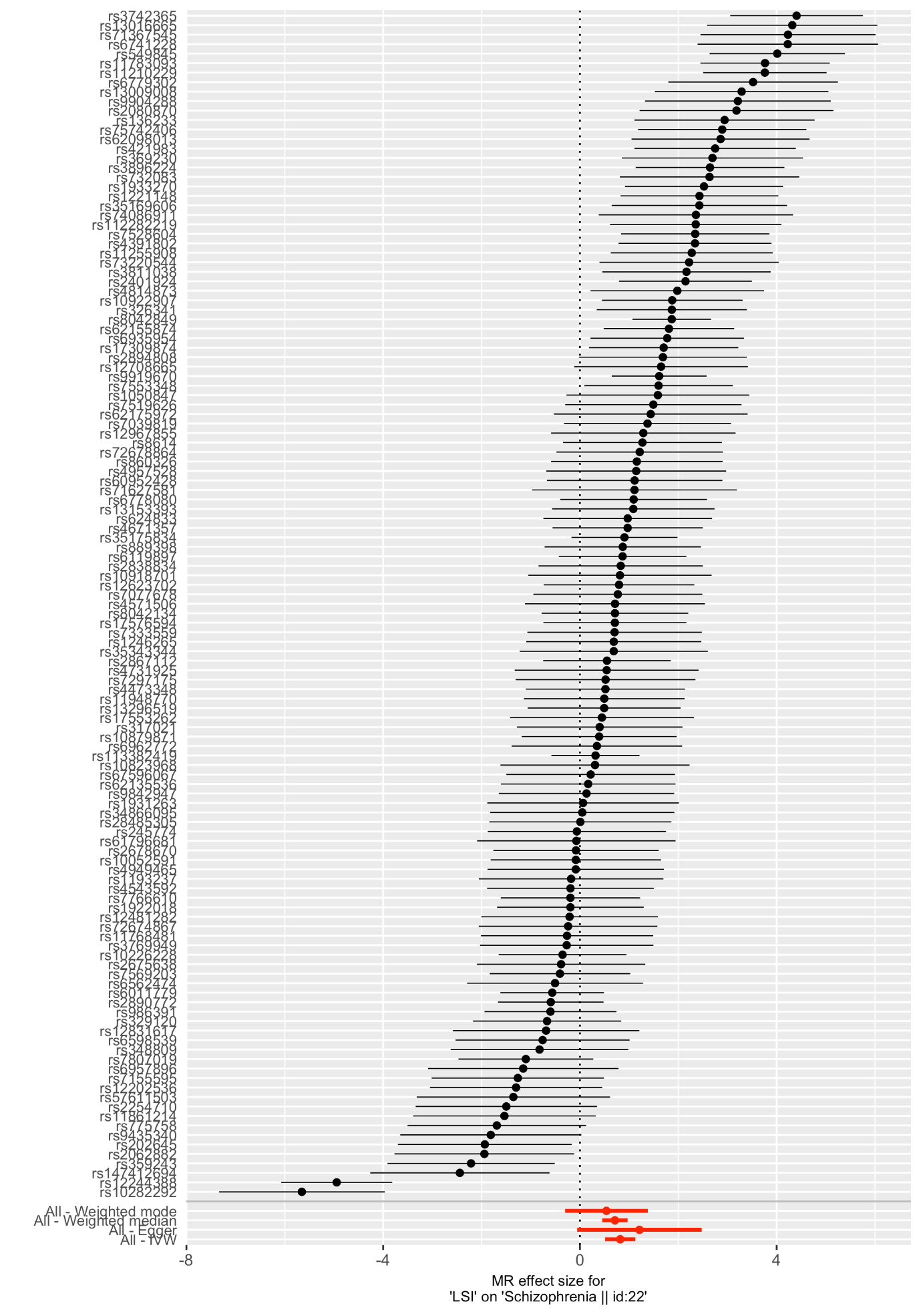

**Figure S13. Single SNP analysis of lifetime smoking on depression.**

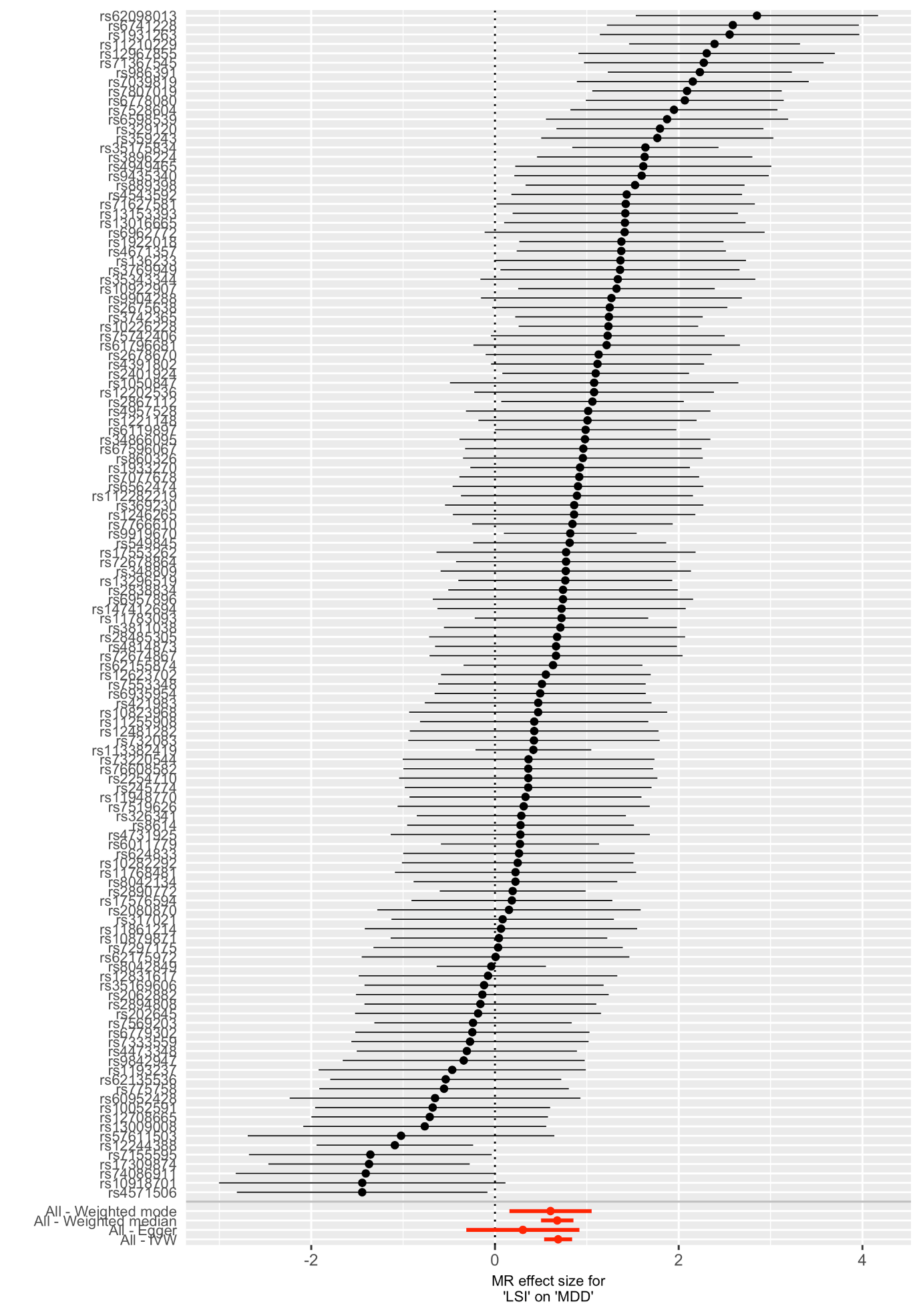

**Figure S14. Single SNP analysis of schizophrenia on lifetime smoking.**

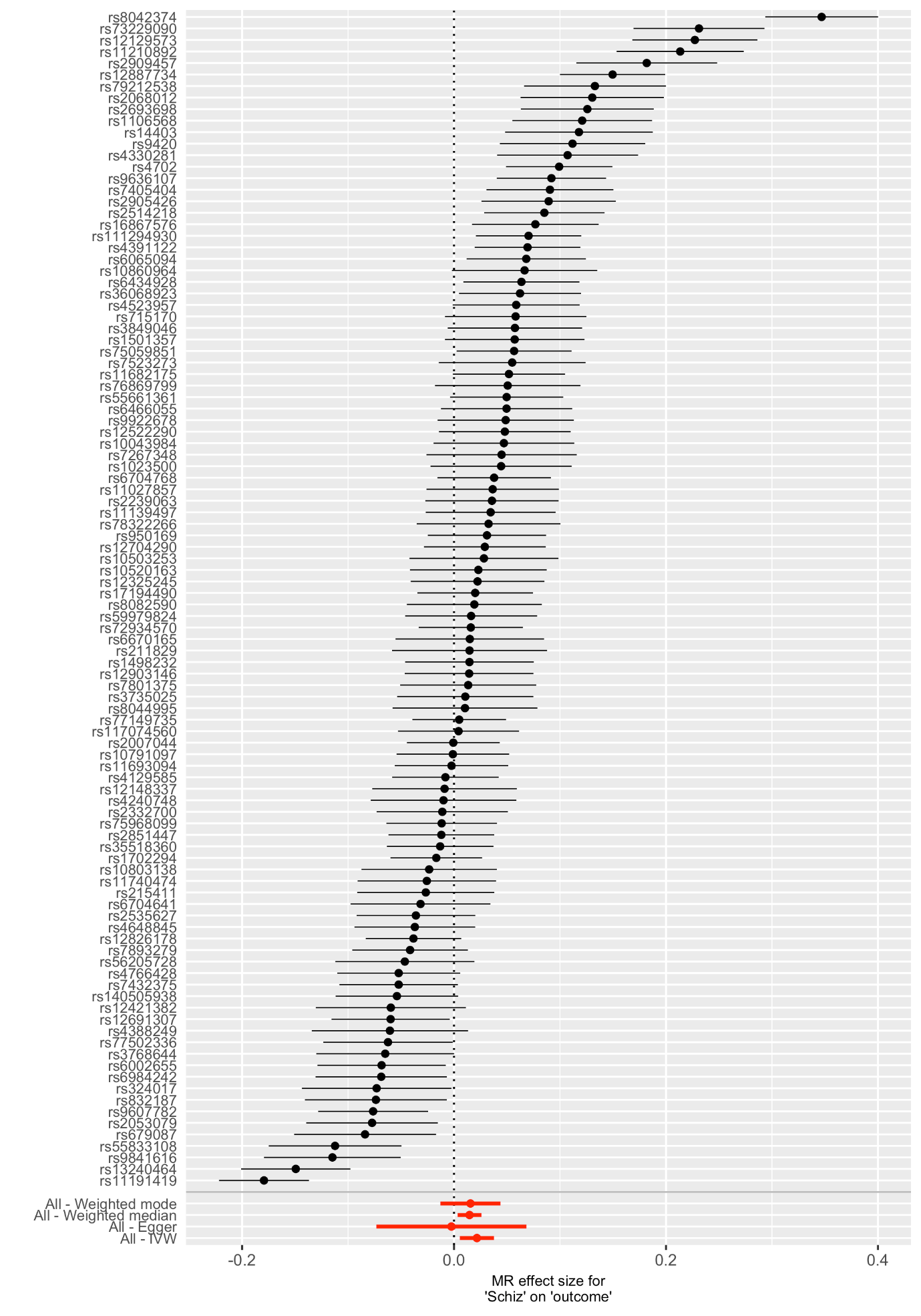

The SNP with the largest effect on smoking is from rs8042374 located in the CHRNA3 gene (known to be associated with Nicotine dependence ^14–17^. However, removing this SNP did not remove the effect (see Supplementary Figure S18).

**Figure S15. Single SNP analysis of depression on lifetime smoking.**

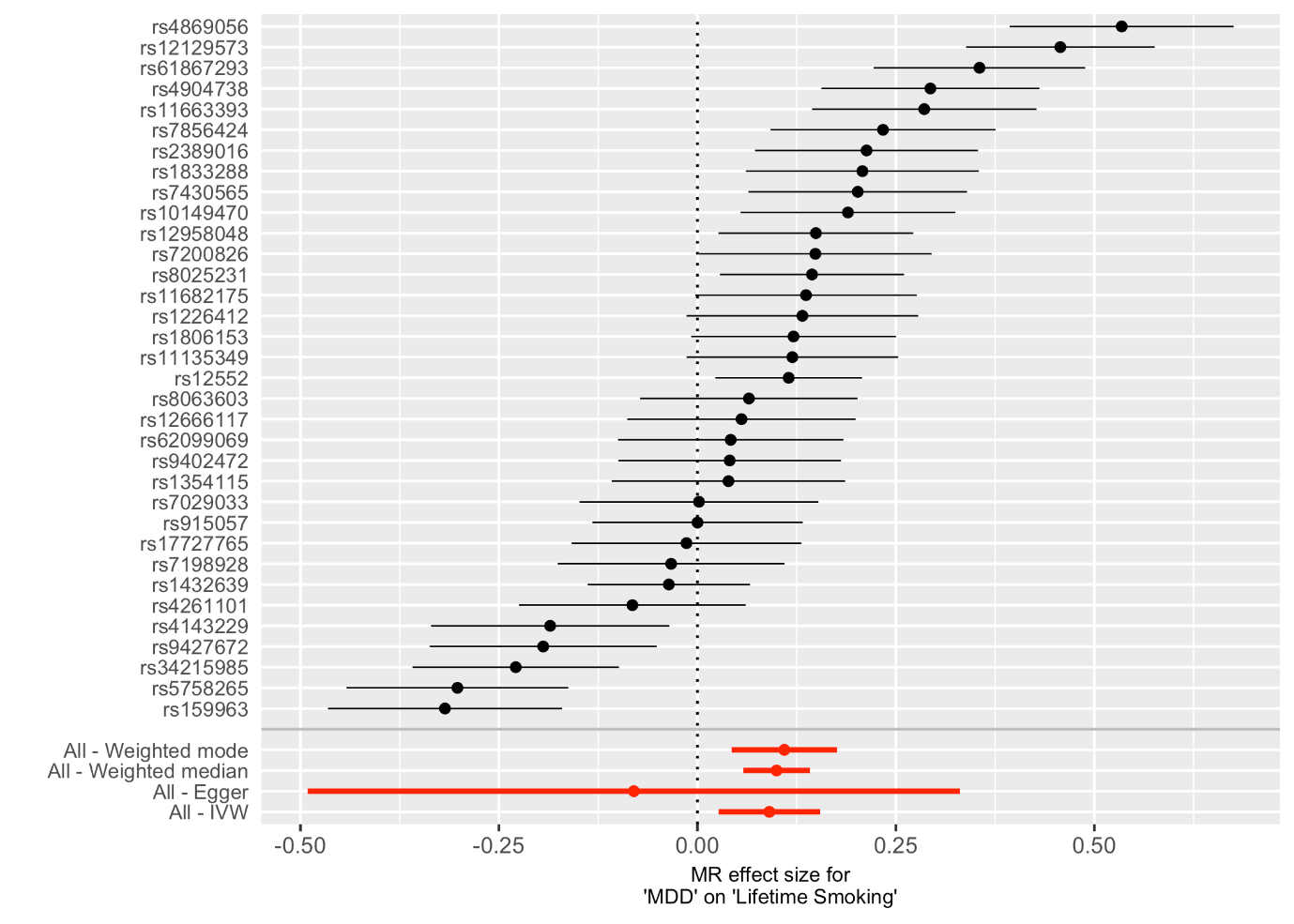

**Figure S16. Leave-one-out analysis of lifetime smoking on schizophrenia.**

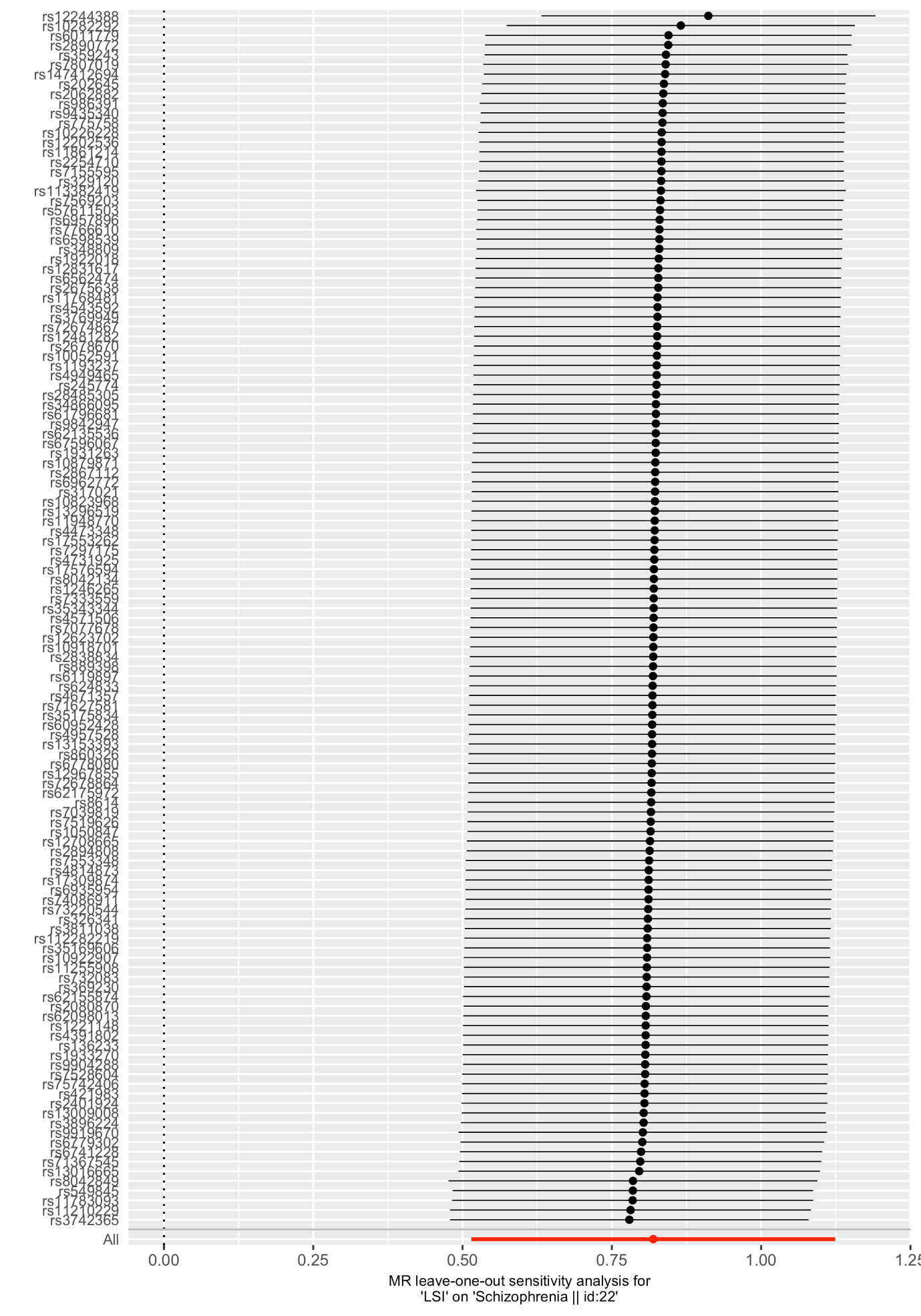

**Figure S17. Leave-one-out analysis of lifetime smoking on depression.**

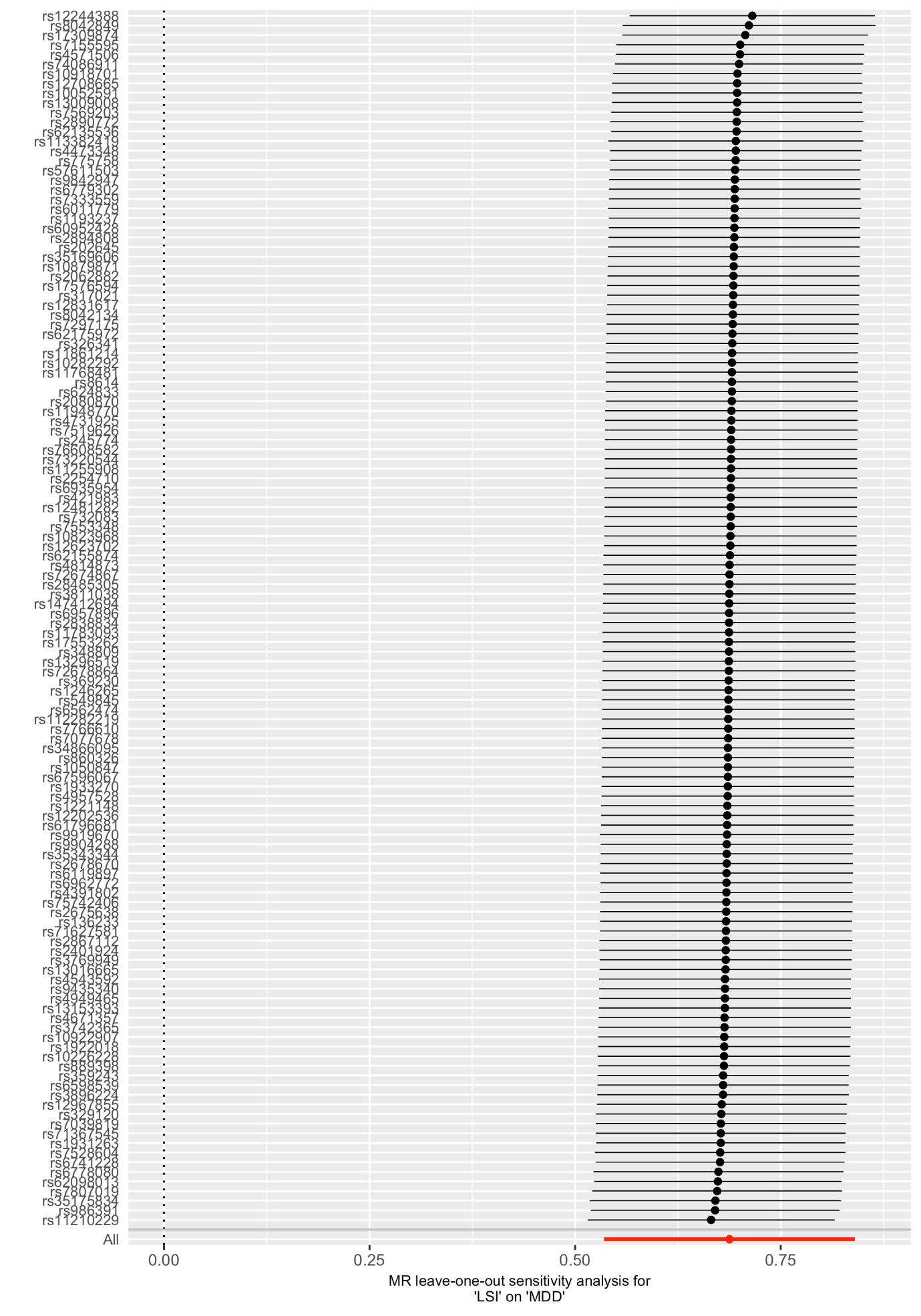

**Figure 18. Leave-one-out analysis of schizophrenia on lifetime smoking.**

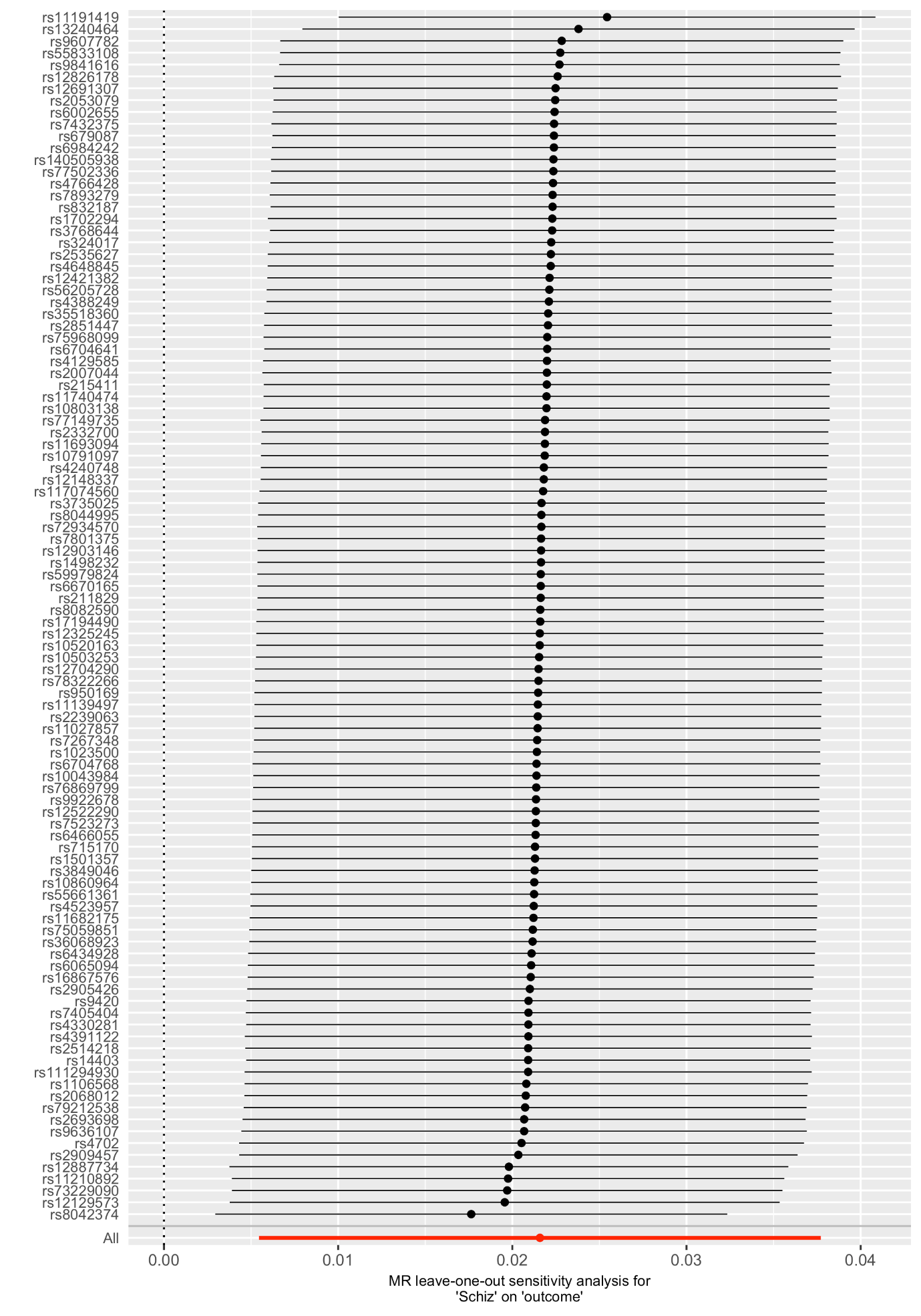

**Figure S19. Leave-one-out analysis of depression on lifetime smoking.**

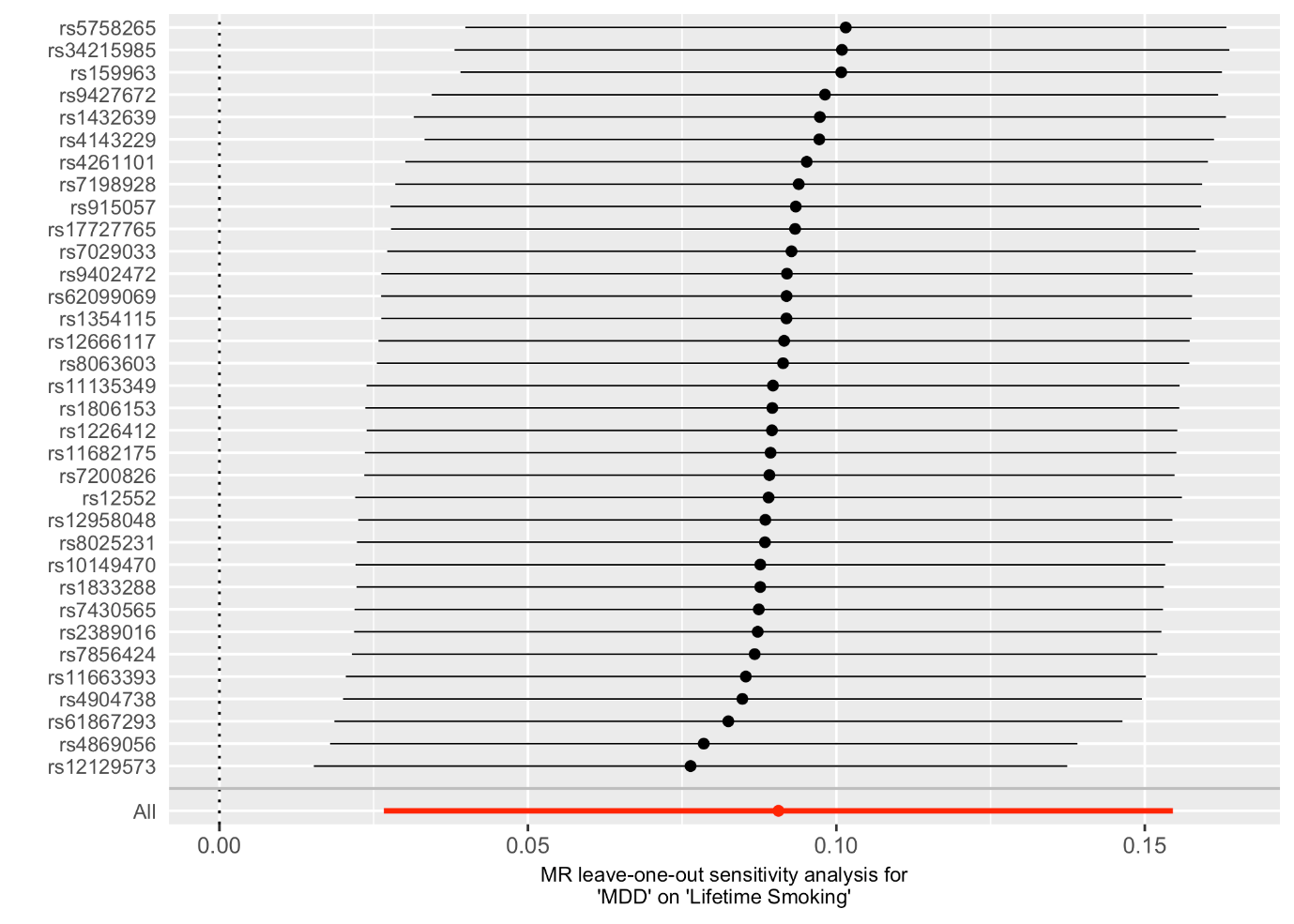

**Table S1. SNPs associated with lifetime smoking index at the genome-wide level of significance (p<5x10^-8^) and clumped for independence at kb=10000 and r^2^=0.001, in ascending order of p-value.**

| **SNP** | **CHR** | **BP** | **EA** | **NonEA** | **EAF** | **Beta** | **SE** | **p-value** |
| --- | --- | --- | --- | --- | --- | --- | --- | --- |
| rs8042849 | 15 | 78817929 | C | T | 0.342 | 0.028 | 0.002 | 1.80E-39 |
| rs113382419 | 9 | 1.36E+08 | C | A | 0.889 | -0.041 | 0.003 | 3.00E-37 |
| rs6011779 | 20 | 61984317 | C | T | 0.191 | 0.028 | 0.003 | 2.30E-27 |
| rs9919670 | 11 | 1.13E+08 | G | A | 0.612 | -0.022 | 0.002 | 7.60E-27 |
| rs2890772 | 2 | 1.46E+08 | G | T | 0.413 | -0.020 | 0.002 | 2.10E-22 |
| rs35175834 | 15 | 47680815 | G | A | 0.788 | -0.024 | 0.002 | 4.60E-22 |
| rs12244388 | 10 | 1.05E+08 | G | A | 0.661 | -0.019 | 0.002 | 1.40E-19 |
| rs11783093 | 8 | 27425349 | C | T | 0.839 | 0.023 | 0.003 | 1.20E-16 |
| rs11210229 | 1 | 73860028 | A | G | 0.384 | 0.017 | 0.002 | 2.00E-16 |
| rs62155874 | 2 | 1.06E+08 | A | G | 0.873 | -0.024 | 0.003 | 5.20E-16 |
| rs10226228 | 7 | 32315613 | A | G | 0.630 | -0.016 | 0.002 | 2.00E-15 |
| rs6119897 | 20 | 31145415 | G | A | 0.762 | -0.018 | 0.002 | 3.60E-15 |
| rs2867112 | 2 | 651349 | T | G | 0.835 | 0.021 | 0.003 | 4.80E-15 |
| rs986391 | 5 | 1.67E+08 | G | A | 0.367 | 0.016 | 0.002 | 9.40E-15 |
| rs3742365 | 14 | 1.04E+08 | T | C | 0.595 | -0.016 | 0.002 | 2.50E-14 |
| rs2401924 | 7 | 1.15E+08 | G | C | 0.502 | 0.015 | 0.002 | 2.70E-14 |
| rs7807019 | 7 | 1.18E+08 | A | G | 0.540 | -0.015 | 0.002 | 6.70E-14 |
| rs549845 | 1 | 44076469 | G | A | 0.301 | 0.016 | 0.002 | 8.30E-14 |
| rs10922907 | 1 | 91193049 | A | T | 0.451 | 0.015 | 0.002 | 3.00E-13 |
| rs7569203 | 2 | 45154418 | A | C | 0.689 | -0.016 | 0.002 | 7.40E-13 |
| rs17309874 | 11 | 27667236 | G | A | 0.740 | -0.016 | 0.002 | 9.70E-13 |
| rs6778080 | 3 | 49317338 | T | C | 0.267 | 0.016 | 0.002 | 1.30E-12 |
| rs8042134 | 15 | 97514404 | T | G | 0.541 | -0.014 | 0.002 | 1.30E-12 |
| rs17576594 | 4 | 1.48E+08 | G | A | 0.724 | 0.016 | 0.002 | 1.70E-12 |
| rs7766610 | 6 | 1.12E+08 | C | A | 0.183 | 0.018 | 0.003 | 2.20E-12 |
| rs1922018 | 7 | 3560401 | C | T | 0.364 | 0.014 | 0.002 | 3.00E-12 |
| rs7553348 | 1 | 75005067 | G | A | 0.438 | 0.014 | 0.002 | 5.20E-12 |
| rs7528604 | 1 | 66407352 | G | A | 0.566 | 0.014 | 0.002 | 5.70E-12 |
| rs329120 | 5 | 1.34E+08 | C | T | 0.581 | 0.014 | 0.002 | 6.30E-12 |
| rs12623702 | 2 | 2.03E+08 | A | G | 0.613 | -0.014 | 0.002 | 7.70E-12 |
| rs13296519 | 9 | 1.28E+08 | G | T | 0.606 | -0.014 | 0.002 | 8.10E-12 |
| rs6935954 | 6 | 26255451 | A | G | 0.421 | 0.014 | 0.002 | 8.20E-12 |
| rs4671357 | 2 | 60136176 | T | C | 0.519 | -0.014 | 0.002 | 1.10E-11 |
| rs3896224 | 10 | 1.06E+08 | A | G | 0.585 | 0.014 | 0.002 | 1.10E-11 |
| rs326341 | 3 | 1.08E+08 | G | A | 0.525 | 0.014 | 0.002 | 1.20E-11 |
| rs4391802 | 11 | 28674592 | A | G | 0.707 | 0.015 | 0.002 | 1.40E-11 |
| rs72678864 | 4 | 1.12E+08 | G | A | 0.829 | 0.018 | 0.003 | 1.60E-11 |
| rs112282219 | 11 | 46632809 | G | A | 0.959 | -0.033 | 0.005 | 3.80E-11 |
| rs10879871 | 12 | 75380511 | T | G | 0.343 | -0.014 | 0.002 | 5.00E-11 |
| rs889398 | 16 | 69556715 | C | T | 0.588 | 0.013 | 0.002 | 6.30E-11 |
| rs4473348 | 2 | 1.82E+08 | A | T | 0.250 | -0.015 | 0.002 | 6.40E-11 |
| rs1221148 | 9 | 1.22E+08 | C | G | 0.587 | 0.013 | 0.002 | 7.30E-11 |
| rs317021 | 4 | 35418368 | T | A | 0.814 | -0.017 | 0.003 | 1.10E-10 |
| rs1933270 | 1 | 49977965 | T | G | 0.364 | 0.013 | 0.002 | 1.50E-10 |
| rs8614 | 17 | 27588806 | C | A | 0.817 | -0.017 | 0.003 | 1.80E-10 |
| rs11255908 | 10 | 8802912 | T | G | 0.743 | -0.015 | 0.002 | 2.30E-10 |
| rs13153393 | 5 | 1.68E+08 | A | G | 0.884 | -0.020 | 0.003 | 2.50E-10 |
| rs2678670 | 2 | 1.04E+08 | A | T | 0.486 | 0.013 | 0.002 | 3.10E-10 |
| rs7333559 | 13 | 1.01E+08 | G | A | 0.212 | 0.015 | 0.002 | 3.20E-10 |
| rs76608582 | 19 | 4474725 | C | A | 0.953 | 0.031 | 0.005 | 3.20E-10 |
| rs421983 | 3 | 84892866 | T | C | 0.519 | 0.013 | 0.002 | 3.30E-10 |
| rs4543592 | 9 | 3014254 | T | C | 0.520 | -0.012 | 0.002 | 4.50E-10 |
| rs11948770 | 5 | 13246336 | T | C | 0.768 | -0.015 | 0.002 | 4.90E-10 |
| rs7039819 | 9 | 82430418 | G | A | 0.427 | 0.013 | 0.002 | 5.10E-10 |
| rs10282292 | 7 | 1.11E+08 | C | T | 0.362 | 0.013 | 0.002 | 5.90E-10 |
| rs2838834 | 21 | 46665208 | C | T | 0.699 | -0.013 | 0.002 | 6.30E-10 |
| rs624833 | 4 | 2881256 | T | G | 0.695 | 0.013 | 0.002 | 6.60E-10 |
| rs62135536 | 2 | 44326028 | C | T | 0.968 | 0.035 | 0.006 | 8.00E-10 |
| rs3811038 | 2 | 1.13E+08 | T | C | 0.724 | -0.014 | 0.002 | 8.90E-10 |
| rs359243 | 2 | 60475509 | T | C | 0.393 | -0.013 | 0.002 | 9.50E-10 |
| rs11768481 | 7 | 96629103 | C | A | 0.666 | 0.013 | 0.002 | 9.90E-10 |
| rs6779302 | 3 | 16859710 | G | T | 0.633 | -0.013 | 0.002 | 1.20E-09 |
| rs35169606 | 8 | 9604066 | T | G | 0.612 | 0.013 | 0.002 | 1.20E-09 |
| rs67596067 | 17 | 50333733 | G | A | 0.649 | -0.013 | 0.002 | 1.20E-09 |
| rs2675638 | 10 | 63576286 | G | A | 0.581 | 0.012 | 0.002 | 1.30E-09 |
| rs75742406 | 11 | 17070365 | G | A | 0.739 | 0.014 | 0.002 | 1.30E-09 |
| rs71367545 | 18 | 77576337 | G | A | 0.791 | -0.015 | 0.002 | 1.40E-09 |
| rs71627581 | 5 | 43161351 | G | A | 0.889 | 0.019 | 0.003 | 1.60E-09 |
| rs13016665 | 2 | 57995348 | C | A | 0.577 | -0.012 | 0.002 | 1.80E-09 |
| rs369230 | 16 | 89645437 | G | T | 0.308 | -0.013 | 0.002 | 1.80E-09 |
| rs10052591 | 5 | 50812738 | T | C | 0.573 | 0.012 | 0.002 | 2.10E-09 |
| rs3769949 | 2 | 1.66E+08 | T | A | 0.528 | -0.012 | 0.002 | 2.50E-09 |
| rs7155595 | 14 | 77502546 | A | C | 0.674 | -0.013 | 0.002 | 2.50E-09 |
| rs7077678 | 10 | 1.04E+08 | C | T | 0.623 | 0.012 | 0.002 | 2.60E-09 |
| rs860326 | 14 | 57342912 | C | T | 0.428 | 0.012 | 0.002 | 2.70E-09 |
| rs12202536 | 6 | 67475273 | A | G | 0.513 | -0.012 | 0.002 | 2.80E-09 |
| rs4814873 | 20 | 19616429 | C | T | 0.767 | 0.014 | 0.002 | 2.90E-09 |
| rs147412694 | 21 | 40702786 | G | A | 0.850 | -0.017 | 0.003 | 2.90E-09 |
| rs9842947 | 3 | 1.57E+08 | C | T | 0.326 | -0.013 | 0.002 | 3.10E-09 |
| rs2894808 | 6 | 52861990 | T | A | 0.922 | -0.022 | 0.004 | 3.50E-09 |
| rs12708665 | 16 | 24728227 | A | G | 0.285 | -0.013 | 0.002 | 3.50E-09 |
| rs202645 | 22 | 41798520 | A | G | 0.203 | -0.015 | 0.002 | 3.90E-09 |
| rs62098013 | 18 | 50863861 | G | A | 0.640 | -0.012 | 0.002 | 4.10E-09 |
| rs4957528 | 5 | 1.06E+08 | A | C | 0.208 | -0.015 | 0.002 | 4.20E-09 |
| rs1246265 | 9 | 86761745 | T | C | 0.305 | -0.013 | 0.002 | 4.20E-09 |
| rs6598539 | 15 | 99204483 | T | C | 0.489 | -0.012 | 0.002 | 4.50E-09 |
| rs13009008 | 2 | 1.74E+08 | A | G | 0.328 | 0.012 | 0.002 | 4.60E-09 |
| rs17553262 | 10 | 92912773 | A | C | 0.885 | -0.018 | 0.003 | 5.30E-09 |
| rs7297175 | 12 | 56473808 | T | C | 0.431 | -0.012 | 0.002 | 6.60E-09 |
| rs245774 | 5 | 1.71E+08 | A | G | 0.272 | -0.013 | 0.002 | 7.40E-09 |
| rs6962772 | 7 | 99081730 | A | G | 0.846 | 0.016 | 0.003 | 7.80E-09 |
| rs12481282 | 20 | 44761377 | G | C | 0.722 | -0.013 | 0.002 | 7.80E-09 |
| rs35343344 | 19 | 18471610 | C | A | 0.733 | 0.013 | 0.002 | 8.80E-09 |
| rs6562474 | 13 | 67332812 | C | G | 0.651 | 0.012 | 0.002 | 1.00E-08 |
| rs775758 | 3 | 77582005 | A | T | 0.433 | 0.012 | 0.002 | 1.10E-08 |
| rs2062882 | 8 | 91839576 | G | A | 0.587 | -0.012 | 0.002 | 1.10E-08 |
| rs7519626 | 1 | 99514554 | C | T | 0.324 | 0.012 | 0.002 | 1.20E-08 |
| rs9435340 | 1 | 1.08E+08 | T | A | 0.344 | 0.012 | 0.002 | 1.20E-08 |
| rs34866095 | 11 | 16377356 | A | G | 0.686 | -0.012 | 0.002 | 1.20E-08 |
| rs348809 | 20 | 59032097 | A | G | 0.348 | -0.012 | 0.002 | 1.30E-08 |
| rs1050847 | 16 | 87443734 | C | T | 0.426 | 0.011 | 0.002 | 1.40E-08 |
| rs73220544 | 3 | 1.31E+08 | A | C | 0.842 | -0.016 | 0.003 | 1.50E-08 |
| rs4571506 | 5 | 87756918 | C | T | 0.540 | 0.011 | 0.002 | 1.50E-08 |
| rs732083 | 17 | 37834367 | G | A | 0.333 | 0.012 | 0.002 | 1.50E-08 |
| rs6741228 | 2 | 22548774 | T | C | 0.433 | 0.011 | 0.002 | 1.60E-08 |
| rs4949465 | 1 | 32178489 | T | C | 0.870 | -0.017 | 0.003 | 1.70E-08 |
| rs62175972 | 2 | 1.61E+08 | T | C | 0.966 | 0.031 | 0.006 | 1.70E-08 |
| rs136233 | 22 | 31212410 | A | G | 0.809 | -0.014 | 0.003 | 1.80E-08 |
| rs12831617 | 12 | 84758368 | C | T | 0.764 | -0.013 | 0.002 | 1.90E-08 |
| rs11861214 | 16 | 746611 | G | T | 0.784 | 0.014 | 0.002 | 2.00E-08 |
| rs10918701 | 1 | 1.62E+08 | G | A | 0.372 | 0.012 | 0.002 | 2.10E-08 |
| rs10823968 | 10 | 74738269 | A | T | 0.633 | 0.012 | 0.002 | 2.10E-08 |
| rs74086911 | 12 | 50015942 | G | A | 0.925 | 0.021 | 0.004 | 2.10E-08 |
| rs4731925 | 7 | 1.33E+08 | C | T | 0.316 | -0.012 | 0.002 | 2.60E-08 |
| rs28485305 | 15 | 74044197 | C | T | 0.631 | 0.012 | 0.002 | 2.60E-08 |
| rs1193237 | 1 | 7526486 | G | C | 0.439 | -0.011 | 0.002 | 2.80E-08 |
| rs60952428 | 16 | 75640521 | T | C | 0.909 | 0.019 | 0.003 | 3.00E-08 |
| rs9904288 | 17 | 47031973 | T | C | 0.708 | 0.012 | 0.002 | 3.10E-08 |
| rs12967855 | 18 | 35138245 | A | G | 0.331 | 0.012 | 0.002 | 3.10E-08 |
| rs2254710 | 6 | 37477000 | C | A | 0.236 | 0.013 | 0.002 | 3.50E-08 |
| rs72674867 | 8 | 95578201 | A | T | 0.765 | 0.013 | 0.002 | 3.80E-08 |
| rs1931263 | 1 | 96175101 | G | T | 0.510 | -0.011 | 0.002 | 4.00E-08 |
| rs57611503 | 16 | 31165795 | G | A | 0.485 | 0.011 | 0.002 | 4.00E-08 |
| rs61796681 | 4 | 23678196 | A | T | 0.912 | -0.019 | 0.004 | 4.20E-08 |
| rs6957896 | 7 | 1.32E+08 | C | T | 0.503 | -0.011 | 0.002 | 4.50E-08 |
| rs2080870 | 5 | 60388313 | A | T | 0.258 | 0.012 | 0.002 | 4.90E-08 |

Note. CHR = chromosome, BP = base position, EA = effect allele, Non EA = non-effect allele, EAF = effect allele frequency.

**Table S2. Tests of the unweighted and weighted regression dilution I^2^_GX._**

|  | I^2^_GX_ Unweighted | I^2^_GX_ Weighted | mF |
| --- | --- | --- | --- |
| Lifetime smoking > CAD | 0.644 | 0.454 | 44.05 |
| Lifetime smoking > Lung cancer | 0.643 | 0.340 | 44.36 |
| Lifetime smoking > AHRR methylation | 0.635 | 0.455 | 44.08 |
| Lifetime smoking > Schizophrenia (2014) | 0.634 | 0.405 | 44.08 |
| Lifetime smoking > Schizophrenia (Steiger filtered) | 0.667 | 0.441 | 44.25 |
| Lifetime smoking > Schizophrenia (2018) | 0.642 | 0.417 | 44.27 |
| Lifetime smoking > Depression (2018) | 0.644 | 0.429 | 44.05 |
| Lifetime smoking > Depression (2018) (Steiger filtered) | 0.646 | 0.436 | 44.22 |
| Lifetime smoking > Depression (2013) | 0 | 0 | 41.06 |
| Schizophrenia (2014) > Lifetime smoking | 0.429 | 0 | 37.90 |
| Schizophrenia (2018) > Lifetime smoking | 0.487 | 0 | 40.86 |
| Depression (2018) > Lifetime smoking | 0 | 0 | 36.72 |
| Depression (2018) > Lifetime smoking (Steiger filtered) | 0 | 0 | 36.49 |
| Depression (2013) > Lifetime smoking | 0.358 | 0 | 19.05 |
| Smoking initiation > Schizophrenia | 0.603 | 0.401 | 44.98 |
| Smoking initiation > Depression (2018) | 0.613 | 0.561 | 44.93 |
| Schizophrenia > Smoking initiation | 0.422 | 0.762 | 37.62 |
| Depression (2018) > Smoking initiation | 0 | 0 | 36.83 |

CAD: coronary artery disease. Unweighted estimates only take into account dilution in the SNP-exposure effects, whereas weighted estimates account for the SE of the SNP-outcome effects^18^. The unweighted I^2^ estimates were larger for both positive control outcomes so in Table 1 (main text), unweighted MR Egger SIMEX estimates are presented. For the main analysis unweighted MR Egger SIMEX corrections are presented unless I^2^ estimates < 0.6 which is too low to conduct either MR Egger analysis. All estimates show evidence of high dilution in the SNP-exposure effects, so MR Egger estimates should be interpreted with caution.

**Table S3. Tests of Heterogeneity in the SNP-exposure association**

|  | Method | Q | df | P-value |
| --- | --- | --- | --- | --- |
| Lifetime smoking > CAD | Inverse-variance weighted | 184.72 | 125 | <0.001 |
|  | MR Egger | 175.12 | 124 | 0.002 |
|  | Q’ | 9.60 | 1 | 0.002 |
| Lifetime smoking > Lung cancer | Inverse-variance weighted | 235.82 | 119 | <0.001 |
|  | MR Egger | 228.96 | 118 | <0.001 |
|  | Q’ | 6.87 | 1 | 0.009 |
| Lifetime smoking > AHRR methylation | Inverse-variance weighted | 122.07 | 118 | 0.38 |
|  | MR Egger | 121.35 | 117 | 0.37 |
|  | Q’ | 0.72 | 1 | 0.40 |
| Lifetime smoking > Schizophrenia | Inverse-variance weighted | 573.03 | 124 | <0.001 |
|  | MR Egger | 571.23 | 123 | <0.001 |
|  | Q’ | 1.81 | 1 | 0.18 |
| Lifetime smoking > Schizophrenia (2018) | Inverse-variance weighted | 592.94 | 121 | <0.001 |
|  | MR Egger | 591.77 | 120 | <0.001 |
|  | Q’ | 1.17 | 1 | 0.28 |
| Lifetime smoking > Depression (2018) | Inverse-variance weighted | 259.98 | 125 | <0.001 |
|  | MR Egger | 256.69 | 124 | <0.001 |
|  | Q’ | 3.29 | 1 | 0.07 |
| Lifetime smoking > Depression (2013) | Inverse-variance weighted | 33.014 | 33 | 0.46 |
|  | MR Egger | 33.07 | 32 | 0.41 |
|  | Q’ | 0.08 | 1 | 0.78 |
| Schizophrenia > Lifetime smoking | Inverse-variance weighted | 773.92 | 101 | <0.001 |
|  | MR Egger | 770.30 | 100 | <0.001 |
|  | Q’ | 3.61 | 1 | 0.05 |
| Schizophrenia (2018) > Lifetime smoking | Inverse-variance weighted | 869.67 | 135 | <0.001 |
|  | MR Egger | 864.25 | 134 | <0.001 |
|  | Q’ | 5.42 | 1 | 0.20 |
| Depression (2018) > Lifetime smoking | Inverse-variance weighted | 255.61 | 33 | <0.001 |
|  | MR Egger | 250.29 | 32 | <0.001 |
|  | Q’ | 5.32 | 1 | 0.02 |
| Depression (2013) > Lifetime smoking | Inverse-variance weighted | 59.75 | 36 | 0.008 |
|  | MR Egger | 59.48 | 35 | 0.006 |
|  | Q’ | 0.27 | 1 | 0.60 |
| Smoking Initiation > Schizophrenia | Inverse-variance weighted | 1468.61 | 370 | <0.001 |
|  | MR Egger | 1467.30 | 369 | <0.001 |
|  | Q’ | 1.32 | 1 | 0.25 |
| Smoking Initiation > Depression (2018) | Inverse-variance weighted | 704.94 | 369 | <0.001 |
|  | MR Egger | 699.86 | 368 | <0.001 |
|  | Q’ | 5.07 | 1 | 0.02 |
| Schizophrenia > Smoking initiation | Inverse-variance weighted | 409.42 | 106 | <0.001 |
|  | MR Egger | 402.35 | 105 | <0.001 |
|  | Q’ | 7.06 | 1 | <0.001 |
| Depression (2018) > Smoking initiation | Inverse-variance weighted | 153.61 | 33 | <0.001 |
|  | MR Egger | 150.95 | 32 | <0.001 |
|  | Q’ | 2.66 | 1 | 0.10 |

Note: df = degrees of freedom where degrees of freedom is equal to the number of SNPs -1. Q = Rucker’s Q^19^, a test of heterogeneity or dispersion in the SNP-exposure effects.

The Q’ indicates the extent to which MR Egger is a better fit than the inverse-variance weighted method.

**Table S4. MR Egger test of directional pleiotropy**

| **Outcome** | **Intercept (95% CI)** | **P-value** |
| --- | --- | --- |
| Lifetime smoking > CAD | 0.012 (0.003, 0.022) | 0.010 |
| Lifetime smoking > Lung cancer | -0.021 (-0.042, -0.001) | 0.062 |
| Lifetime smoking > AHRR methylation | -0.001 (-0.004, 0.002) | 0.406 |
| Lifetime smoking > Schizophrenia (2014) | -0.006 (-0.025, 0.013) | 0.531 |
| Lifetime smoking > Schizophrenia (2018) | -0.004 (-0.022, 0.013) | 0.627 |
| Lifetime smoking > Depression (2018) | 0.006 (-0.003, 0.015) | 0.210 |
| Lifetime smoking > Depression (2013) | 0.009 (-0.055, 0.073) | 0.787 |
| Schizophrenia (2014) > Lifetime smoking | 0.002 (-0.004, 0.007) | 0.495 |
| Schizophrenia (2018) > Lifetime smoking | 0.002 (-0.002, 0.006) | 0.360 |
| Depression (2018) > Lifetime smoking | 0.006 (-0.008, 0.019) | 0.416 |
| Depression (2013) > Lifetime smoking | -0.001 (-0.004, 0.003) | 0.693 |
| Smoking initiation > Schizophrenia | -0.003 (-0.013, 0.007) | 0.565 |
| Smoking initiation > Depression (2018) | 0.004 (-0,001, 0.010) | 0.103 |
| Schizophrenia > Smoking initiation | 0.002 (-0.001, 0.006) | 0.177 |
| Depression (2018) > Smoking initiation | 0.003 (-0.012, 0.006) | 0.458 |

CAD: coronary artery disease.

**Table S5. Bi-directional two-sample Mendelian randomisation of lifetime smoking on schizophrenia and depression following Steiger filtering.**

| **Exposure** | **Outcome** | **Method** | **N SNP** | **OR (95% CI)** | **p-value** |
| --- | --- | --- | --- | --- | --- |
| **Smoking** | **Schizophrenia** | Inverse-Variance Weighted | 105/125 (84%) | 1.73 (1.39, 2.16) | 9.12 x 10^-07^ |
|  |  | MR Egger (SIMEX) |  | 6.19 (2.75, 13.95) | 2.64 x 10^-05^ |
|  |  | Weighted median |  | 1.72 (1.34, 2.22) | 2.78 x 10^-05^ |
|  |  | Weighted mode |  | 1.84 (0.73, 4.65) | 0.20 |
|  |  | MR RAPS |  | 1.83 (1.45, 2.32) | 3.97 x 10^-07^ |
| **Smoking** | **Depression** | Inverse-Variance Weighted | 124/126 (98%) | 1.94 (1.67, 2.25) | 3.46 x 10^-18^ |
|  |  | MR Egger (SIMEX) |  | 1.17 (0.67, 2.04) | 0.59 |
|  |  | Weighted median |  | 1.95 (1.65, 2.31) | 4.70 x 10^-15^ |
|  |  | Weighted mode |  | 1.81 (1.17, 2.79) | 0.008 |
|  |  | MR RAPS |  | 1.95 (1.68, 2.27) | 3.81 x 10^-18^ |
| **Exposure** | **Outcome** | **Method** | **N SNP** | **Beta (95% CI)** | **p-value** |
| **Schizophrenia** | **Smoking** | - | 102/102 (100%) | - | - |
| **Depression** | **Smoking** | Inverse-Variance Weighted | 32/34 (94%) | 0.063 (0.007, 0.120) | 0.028 |
|  |  | MR Egger (SIMEX) |  | - | - |
|  |  | Weighted median |  | 0.084 (0.042, 0.126) | 8.64 x 10^-05^ |
|  |  | Weighted mode |  | 0.111 (0.034, 0.187) | 0.008 |
|  |  | MR RAPS |  | 0.065 (0.008, 0.123) | 0.026 |

Note: For each SNP in the instrument, Steiger filtering calculates how much of the variance the SNP explains in the exposure and how much it explains in the outcome. The number of SNPs which explain more variance in the exposure are presented in the N SNP column. Analysis is then repeated using only these SNPs to ensure that results are not due to reverse causation. For the effect of schizophrenia on lifetime smoking, all SNPs comprising the instrument for schizophrenia were better instruments for schizophrenia than lifetime smoking, therefore, that analysis was not repeated. All SIMEX corrections are unweighted due to greater unweighted I^2^_GX_ (see Supplementary Table S2). Given low I^2^_GX_, all MR Egger results should be interpreted with caution.

**Table S6. Bi-directional two-sample Mendelian randomisation analyses of the effect of lifetime smoking on depression (2013).**

| **Exposure** | **Outcome** | **Method** | **OR (95% CI)** | **p-value** |
| --- | --- | --- | --- | --- |
| **Smoking** | **Depression** | Inverse-Variance Weighted | 3.66 (2.08, 6.43) | 6.46 x 10^-06^ |
|  |  | Weighted median | 3.69 (1.72, 7.94) | 0.001 |
|  |  | Weighted mode | 3.18 (0.63, 16.06) | 0.17 |
|  |  | MR RAPS | 4.23 (2.33, 7.67) | 2.04 x 10^-06^ |
| **Exposure** | **Outcome** | **Method** | **Beta (95% CI)** | **p-value** |
| **Depression** | **Smoking** | Inverse-Variance Weighted | 0.002 (-0.006, 0.011) | 0.572 |
|  |  | Weighted median | 0.003 (-0.007, 0.012) | 0.602 |
|  |  | Weighted mode | 0.002 (-0.016, 0.019) | 0.844 |
|  |  | MR RAPS | 0.003 (-0.005, 0.012) | 0.435 |

Note: When depression is the exposure, a relaxed p-value threshold of p<5x10^-5^ was used because there were no SNPs associated at the genome wide level of significance. The direction of effect is consistent with the more recent GWAS for MDD despite sample overlap. There is weaker statistical evidence, possibly due to reduced sample size (N = 18 759). Both MR Egger and MR Egger (SIMEX) estimates could not be conducted due to low regression dilution I^2^_GX_ (see Table S2).

**Table S7. Bidirectional two-sample Mendelian randomisation analyses of the effect of lifetime smoking (from a sensitivity GWAS without chip as a covariate) on schizophrenia and major depression.**

| **Exposure** | **Outcome** | **Method** | **N SNP** | **OR (95% CI)** | **P-value** |
| --- | --- | --- | --- | --- | --- |
| **Smoking** | **Schizophrenia** | Inverse-Variance Weighted | 137 | 2.23 (1.69, 2.95) | 2.03 x 10^-08^ |
|  |  | MR Egger (SIMEX) | 137 | 3.71 (1.47, 3.39) | 0.006 |
|  |  | Weighted median | 137 | 2.07 (1.61, 2.66) | 1.24 x 10^-08^ |
|  |  | Weighted mode | 137 | 1.56 (0.63, 3.86) | 0.34 |
|  |  | MR RAPS | 137 | 2.42 (1.86, 3.15) | 6.09 x 10^-11^ |
| **Smoking** | **Depression** | Inverse-Variance Weighted | 90 | 1.88 (1.57, 2.24) | 5.26 x 10^-12^ |
|  |  | MR Egger (SIMEX) | 90 | 4.39 (0.60, 323.15) | 0.51 |
|  |  | Weighted median | 90 | 1.70 (1.39, 2.07) | 1.53 x 10^-07^ |
|  |  | Weighted mode | 90 | 1.49 (0.99, 2.26) | 0.06 |
|  |  | MR RAPS | 90 | 1.85 (1.55, 2.21) | 9.00 x 10^-12^ |
|  |  |  |  | **Beta (95% CI)** | **P-value** |
| **Schizophrenia** | **Smoking** | Inverse-Variance Weighted | 102 | 0.020 (0.003, 0.037) | 0.02 |
|  |  | MR Egger (SIMEX) | 102 | - | - |
|  |  | Weighted median | 102 | 0.008 (-0.004, 0.020) | 0.19 |
|  |  | Weighted Mode | 102 | 0.004 (-0.027, 0.035) | 0.80 |
|  |  | MR RAPS | 102 | 0.014 (-0.001, 0.030) | 0.06 |
| **Depression** | **Smoking** | Inverse-Variance Weighted | 36 | 0.102 (0.036, 0.168) | 0.002 |
|  |  | MR Egger (SIMEX) | - | - | - |
|  |  | Weighted median | 36 | 0.101 (0.056, 0.147) | 1.29 x 10^-05^ |
|  |  | Weighted mode | 36 | 0.124 (0.044, 0.204) | 0.004 |
|  |  | MR RAPS | 36 | 0.091 (0.025, 0.157) | 0.007 |

Note: There were 139 genome-wide significant SNPs associated with lifetime smoking index when genotype chip was not included as a covariate in the GWAS. SIMEX-corrected estimates are unweighted. MR Egger regression was not conducted for schizophrenia or major depression as exposures because regression dilution I^2^_GX_ was below 0.3 (see Supplementary Table S6). Due to low regression dilution I^2^_GX_ for lifetime smoking as the exposure (see Supplementary Table S6), MR Egger and MR Egger SIMEX estimates should be interpreted with caution. SIMEX = simulation extrapolation, MR RAPS = robust adjusted profile score.

**Table S8. Bidirectional two-sample Mendelian randomisation analyses of the effect of smoking initiation on schizophrenia and major depression.**

| **Exposure** | **Outcome** | **Method** | **N SNP** | **OR (95% CI)** | **P-value** |
| --- | --- | --- | --- | --- | --- |
| **Smoking initiation** | **Schizophrenia** | Inverse-Variance Weighted | 371 | 1.53 (1.35, 1.74) | 3.70 x 10^-11^ |
|  |  | MR Egger (SIMEX) | 371 | 1.35 (0.83, 2.22) | 0.23 |
|  |  | Weighted median | 371 | 1.38 (1.23, 1.55) | 3.44 x 10^-08^ |
|  |  | Weighted mode | 371 | 1.23 (0.74, 2.06) | 0.42 |
|  |  | MR RAPS | 371 | 1.63 (1.45, 1.85) | 4.46 x 10^-15^ |
| **Smoking initiation** | **Depression** | Inverse-Variance Weighted | 370 | 1.54 (1.44, 1.64) | 3.61 x 10^-37^ |
|  |  | MR Egger (SIMEX) | 370 | 1.36 (1.10, 1.67) | 0.004 |
|  |  | Weighted median | 370 | 1.46 (1.35, 1.58) | 4.62 x 10^-21^ |
|  |  | Weighted mode | 370 | 1.44 (1.10, 1.89) | 0.008 |
|  |  | MR RAPS | 370 | 1.54 (1.44, 1.65) | 1.31 x 10^-35^ |
|  |  |  |  | **Beta (95% CI)** | **P-value** |
| **Schizophrenia** | **Smoking initiation** | Inverse-Variance Weighted | 107 | 0.010 (0.000, 0.021) | 0.04 |
|  |  | MR Egger (SIMEX) | 107 | -0.030 (-0.093, 0.033) | 0.35 |
|  |  | Weighted median | 107 | 0.003 (-0.006, 0.012) | 0.53 |
|  |  | Weighted Mode | 107 | -0.008 (-0.033, 0.017) | 0.54 |
|  |  | MR RAPS | 107 | 0.008 (-0.002, 0.017) | 0.11 |
| **Depression** | **Smoking initiation** | Inverse-Variance Weighted | 34 | 0.083 (0.039, 0.127) | 2.32 x 10^-04^ |
|  |  | MR Egger (SIMEX) | 34 | - | - |
|  |  | Weighted median | 34 | 0.077 (0.042, 0.112) | 1.55 x 10^-05^ |
|  |  | Weighted mode | 34 | 0.062 (0.007, 0.117) | 0.03 |
|  |  | MR RAPS | 34 | 0.077 (0. 037, 0.117) | 1.64 x 10^-04^ |

Note: Due to low regression dilution I^2^_GX_ (see Supplementary Table S6), MR Egger estimates could not be conducted. For lifetime smoking as the exposure, unweighted MR Egger SIMEX estimates were conducted. For schizophrenia as the exposure, weighted MR Egger SIMEX was conducted. When major depression was the exposure, both weighted and unweighted I^2^_GX_ values were too low for SIMEX to be conducted. SIMEX = simulation extrapolation, MR RAPS = robust adjusted profile score. Smoking initiation scores are given in betas by GSCAN calculated from the meta-analysed z-score statistic by assuming a prevalence for the binary trait^20^.

**Table S9. Bi-directional two-sample Mendelian randomisation analyses of the effect of lifetime smoking on schizophrenia (2018).**

| **Exposure** | **Outcome** | **Method** | **OR (95% CI)** | **p-value** |
| --- | --- | --- | --- | --- |
| **Smoking** | **Schizophrenia** | Inverse-Variance Weighted | 2.64 (1.99, 3.52) | 2.36 x 10^-11^ |
|  |  | MR Egger (SIMEX) | 4.03 (1.38, 11.72) | 0.01 |
|  |  | Weighted median | 2.23 (1.76, 2.82) | 4.02 x 10^-11^ |
|  |  | Weighted mode | 2.20 (1.11, 4.36) | 0.02 |
|  |  | MR RAPS | 2.74 (2.09, 3.58) | 2.55 x 10^-13^ |
| **Exposure** | **Outcome** | **Method** | **Beta (95% CI)** | **p-value** |
| **Schizophrenia** | **Smoking** | Inverse-Variance Weighted | 0.025 (0.011, 0.038) | 0.0003 |
|  |  | MR Egger (SIMEX) | - | - |
|  |  | Weighted median | 0.016 (0.005, 0.026) | 0.004 |
|  |  | Weighted mode | 0.029 (-0.011, 0.068) | 0.159 |
|  |  | MR RAPS | 0.017 (0.004, 0.03) | 0.009 |

Note: MR Egger estimates could not be conducted due to low regression dilution I^2^_GX_ (see Table S2).

**Table S10. A comparison of each of the MR sensitivity analyses and their assumptions.**

| **Method** | **Description** | **Additional Assumptions** | **Power** | **Invalid variants allowed** | **IV2** | **IV3** |
| --- | --- | --- | --- | --- | --- | --- |
| Random effects Inverse‐variance weighted (IVW) | A meta-analysis of the Wald ratios for each SNP ($\frac{ZY}{ZX}$) weighted by the inverse of the variance of the SNP-outcome association. | Any horizontal pleiotropy must be balanced | Has the most power if the assumptions are satisfied. | 0% (or 100% if all horizontal pleiotropy is balanced) | ✗ | ✗ |
| MR‐Egger regression^21^ | An extension of the IVW which relaxes the assumption that any pleiotropy must be balanced. A significant intercept term suggests bias from directional pleiotropy, i.e. the average pleiotropic effect is not zero. MR-Egger regression provides consistent estimates even if all genetic instrumental variables are invalid as long as the INSIDE assumption is met. | InSIDE^a^  NOME^b^ | Has the lowest power | 100% | ✗ | ✓ |
| Weighted median^22^ | The weighted median estimate is obtained by first calculating the Wald ratio causal estimate for each SNP and then taking the estimate with the median inverse variance weight. | Consistent when 50% of weight contributed by genetic variants is valid. | Similar to that of IVW method. | 50% | ✓ | ✓ |
| Weighted mode^23^ | Finds the largest cluster of Wald ratio estimates. The majority of the genetic instruments can be invalid providing the ZEMPA assumption is satisfied. In the weighted mode method, the mode is calculated using the inverse variance weights of the Wald ratios. | ZEMPA^c^ | Less powerful than IVW and weighted median. | 50% | ✓ | ✓ |
| Robust adjusted profile score (RAPS)^24^ | An extension of IVW into a general framework which allows very many weak instruments. Requires no sample overlap in the exposure and outcome SNP effect estimates. | InSIDE^a^  Pleiotropy is additive.  Pleiotropic effects are balanced (have mean zero). |  | 100% | ✓ | ✓ |

Note. Where Z = the genetic instrument, X = the exposure and Y = the outcome. IV2 = assumption 2, that all instruments (Z) must not be associated with confounders. IV3 = assumption 3, that all instruments (Z) must only be associated with the outcome (Y) through the exposure (X). These two columns have a cross if that method requires the assumption to be met and a tick if that assumption can be relaxed. Throughout the table, invalid refers to instruments that do not meet the required assumptions for MR. Power of the methods might differ under different models of pleiotropy. ^a^The InSIDE assumption = pleiotropic effects of Z are independent of the effects of Z on the exposure. ^b^The no measurement error assumption (NOME) = assumes that the ZX associations are known, rather than estimated. ^c^The ZEMPA assumption = the largest subset of genetic instrumental variables with the same ratio estimate will contain the valid instruments.

**References**

1.	Leffondré, K., Abrahamowicz, M., Siemiatycki, J. & Rachet, B. Modeling smoking history: a comparison of different approaches. *Am. J. Epidemiol.* **156**, 813–823 (2002).

2.	Lewer, D., McKee, M., Gasparrini, A., Reeves, A. & de Oliveira, C. Socioeconomic position and mortality risk of smoking: evidence from the English Longitudinal Study of Ageing (ELSA). *Eur. J. Public Health* **27**, 1068–1073 (2017).

3.	Bycroft, C. *et al.* Genome-wide genetic data on ~500,000 UK Biobank participants. *bioRxiv* 166298 (2017). doi:10.1101/166298

4.	Mitchell, R., Hemani, G., Dudding, T. & Paternoster, L. UK Biobank Genetic Data: MRC-IEU Quality Control, Version 1. *data.bris* (2017). doi:10.5523/bris.3074krb6t2frj29yh2b03x3wxj

5.	Boyd, A. *et al.* Cohort profile: the ‘children of the 90s’—the index offspring of the Avon Longitudinal Study of Parents and Children. *Int. J. Epidemiol.* **42**, 111–127 (2013).

6.	Golding, J., Pembrey, M., Jones, R. & ALSPAC Study Team. ALSPAC–the avon longitudinal study of parents and children. *Paediatr. Perinat. Epidemiol.* **15**, 74–87 (2001).

7.	Wang, Y. *et al.* Rare variants of large effect in BRCA2 and CHEK2 affect risk of lung cancer. *Nat. Genet.* **46**, 736 (2014).

8.	CARDIoGRAMplusC4D Consortium & others. A comprehensive 1000 Genomes-based genome-wide association meta-analysis of coronary artery disease. *Nat. Genet.* **47**, 1121–1149 (2015).

9.	Relton, C. L. *et al.* Data resource profile: accessible resource for integrated epigenomic studies (ARIES). *Int. J. Epidemiol.* **44**, 1181–1190 (2015).

10.	Pidsley, R. *et al.* A data-driven approach to preprocessing Illumina 450K methylation array data. *BMC Genomics* **14**, 293 (2013).

11.	Touleimat, N. & Tost, J. Complete pipeline for Infinium® Human Methylation 450K BeadChip data processing using subset quantile normalization for accurate DNA methylation estimation. *Epigenomics* **4**, 325–341 (2012).

12.	Gaunt, T. R. *et al.* Systematic identification of genetic influences on methylation across the human life course. *Genome Biol.* **17**, 61 (2016).

13.	Houseman, E. A. *et al.* DNA methylation arrays as surrogate measures of cell mixture distribution. *BMC Bioinformatics* **13**, 86 (2012).

14.	Munafò, M. R. *et al.* Association Between Genetic Variants on Chromosome 15q25 Locus and Objective Measures of Tobacco Exposure. *JNCI J. Natl. Cancer Inst.* **104**, 740–748 (2012).

15.	Thorgeirsson, T. E. *et al.* A Variant Associated with Nicotine Dependence, Lung Cancer and Peripheral Arterial Disease. *Nature* **452**, 638–642 (2008).

16.	Tobacco Consortium. Genome-wide meta-analyses identify multiple loci associated with smoking behavior. *Nat. Genet.* **42**, 441–447 (2010).

17.	Ware, J. J., van den Bree, M. B. & Munafò, M. R. Association of the CHRNA5-A3-B4 gene cluster with heaviness of smoking: a meta-analysis. *Nicotine Tob. Res.* **13**, 1167–1175 (2011).

18.	Bowden, J. *et al.* Assessing the suitability of summary data for two-sample Mendelian randomization analyses using MR-Egger regression: the role of the I 2 statistic. *Int. J. Epidemiol.* **45**, 1961–1974 (2016).

19.	Bowden, J. *et al.* A framework for the investigation of pleiotropy in two-sample summary data Mendelian randomization. *Stat. Med.* **36**, 1783–1802 (2017).

20.	Liu, M. Association studies of up to 1.2 million individuals yield new insights in the genetic etiology of tobacco and alcohol use. *Nat. Genet.* (In Press).

21.	Bowden, J., Davey Smith, G. & Burgess, S. Mendelian randomization with invalid instruments: effect estimation and bias detection through Egger regression. *Int. J. Epidemiol.* **44**, 512–525 (2015).

22.	Bowden, J., Davey Smith, G., Haycock, P. C. & Burgess, S. Consistent estimation in Mendelian randomization with some invalid instruments using a weighted median estimator. *Genet. Epidemiol.* **40**, 304–314 (2016).

23.	Hartwig, F. P., Smith, G. D. & Bowden, J. Robust inference in two-sample Mendelian randomisation via the zero modal pleiotropy assumption. *bioRxiv* 126102 (2017).

24.	Zhao, Q., Wang, J., Hemani, G., Bowden, J. & Small, D. S. Statistical inference in two-sample summary-data Mendelian randomization using robust adjusted profile score. *arXiv* (2018).
